# *Saccharomyces cerevisiae* DJ-1 paralogs maintain genome integrity through glycation repair of nucleic acids and proteins

**DOI:** 10.1101/2022.09.23.509147

**Authors:** Gautam Susarla, Priyanka Kataria, Amrita Kundu, Patrick D’Silva

## Abstract

Reactive carbonyl species (RCS) such as methylglyoxal and glyoxal are potent glycolytic intermediates that extensively damage cellular biomolecules leading to genetic aberration and protein misfolding. Hence, RCS levels are crucial indicators in the progression of various pathological diseases. Besides the glyoxalase system, emerging studies report highly conserved DJ-1/ThiJ/PfpI superfamily proteins as critical regulators of RCS. DJ-1 superfamily proteins, including the human DJ-1, a genetic determinant of Parkinson’s disease possess diverse physiological functions paramount for combating multiple stressors. Although *S. cerevisiae* retains four DJ-1 orthologs (namely Hsp31, Hsp32, Hsp33, and Hsp34), their physiological relevance and collective requirement are still obscure due to their close sequence similarity. Here, we report for the first time that the yeast DJ-1 orthologs function as novel enzymes involved in the preferential scavenge of glyoxal and methylglyoxal, toxic metabolites, and genotoxic agents. At the cellular level, their collective loss induces chronic glycation of the proteome, and nucleic acids, resulting in a spectrum of genetic mutations and reduced mRNA translational efficiency. Furthermore, the Hsp31 paralogs efficiently repair severely glycated macromolecules derived from carbonyl modifications. They also participate in genome maintenance as their absence upregulates DNA damage response pathways when exposed to different genotoxins. Interestingly, yeast DJ-1 orthologs provide robust organellar protection by redistributing into mitochondria to alleviate the glycation damage of mitochondrial DNA and proteins. Taken together, our study uncovers the existence of a novel glycation repair pathway in *S. cerevisiae* and a possible neuroprotective mechanism of how hDJ-1 confers mitochondrial health during carbonyl stress.

## Introduction

The structural and functional integrity of nucleic acids and proteins is critical for establishing a physiological balance to maintain normal cellular health. Reactive carbonyl species (RCS) are highly toxic compounds that covalently bind and damage vital cellular components such as protein, DNA, and fatty acids (Semchyshyn, 2014). RCS primarily constitutes methylglyoxal (MG), glyoxal (GO), and 3-deoxyglucosone (Semchyshyn, 2014). The MG and GO are metabolic by-products of reactive aldehyde and ketones, respectively (Allaman, Bélanger, & Magistretti, 2015; Lange, Wood, Knight, Assimos, & Holmes, 2012). At the basal physiological level, these dicarbonyls participate in numerous signaling processes essential for cellular development and stress response (Akhand et al., 2001; Rodrigues et al., 2020). On the contrary, at elevated levels, MG and GO modifies the thiol and amine groups through the Maillard reaction forming a myriad of irreversible intermediates followed by stable Advanced glycation end-products (AGEs) (Semchyshyn, 2014; Thornalley, 2008). They preferentially form adducts with side chains of arginine, lysine, and cysteine and indiscriminately impair the structure and function of proteins. Moreover, the glycation of essential proteins, including antioxidant machinery, can alter reactive oxygen species (ROS) homeostasis, mitochondrial performance, and proteostasis (Goudarzi, Kalantari, & Rezaei, 2018; Seo, Ki, & Shin, 2014). Likewise, chronic exposure of nucleic acids to RCS culminates into nucleotide AGEs that promote genetic aberrations and translational defects and induce a spectrum of mutations, including transition and G: C to T: A transversions (Murata-Kamiya, Kamiya, Kaji, & Kasai, 1997, 2000). Elevated glycation of protein and DNA has implications in several pathologies like diabetes, aging, and neurodegenerative diseases (Fournet, Bonté, & Desmoulière, 2018). Therefore, regulating endogenous RCS and minimizing its detrimental effects on biomolecules is critically important. The glyoxalase system that constitutes glyoxalase I and II efficiently detoxifies MG and GO to D-lactate and glycolate separately by utilizing glutathione (GSH) as a cofactor (Inoue, Maeta, & Nomura, 2011). Since excess levels of carbonyls concomitantly induce ROS, the GSH pool is rapidly depleted during the anti-oxidation process (de Bari et al., 2020). Hence, cells have evolved GSH independent pathway (glyoxalase III system) to maintain healthy redox balance, primarily comprising DJ-1/ThiJ/Pfp1 superfamily proteins (Bankapalli et al., 2015; J. Y. Lee et al., 2012; Melvin, Bankapalli, D’Silva, & Shivaprasad, 2017).

DJ-1 superfamily members are highly conserved, multi-stress responding, and ubiquitously present in most organisms (Bandyopadhyay & Cookson, 2004; Wei, Ringe, Wilson, & Ondrechen, 2007). The superfamily includes human DJ-1(hDJ-1/*PARK7)*, a well-explored protein that induces a familial form of Parkinson’s disease (PD) through its genetic mutations (Bonifati et al., 2003). Besides having various neuroprotective functions, hDJ-1 possesses both methylglyoxalase and glyoxalase activity that substantially modulates endogenous carbonyls (J. Y. Lee et al., 2012). Moreover, the deletion of *E. Coli* DJ-1 members exhibited alterations in the glycation of DNA and proteins (C. Lee, Lee, Lee, & Park, 2016; Richarme et al., 2017). *Saccharomyces cerevisiae* has four homologs of DJ-1, namely Hsp31, Hsp32, Hsp33, and Hsp34, representing Hsp31 mini-family proteins (Wilson, St Amour, Collins, Ringe, & Petsko, 2004). Interestingly, Hsp32, Hsp33, and Hsp34 share ∼99.5% sequence identity within them and ∼70% sequence identity with Hsp31. All the paralogs possess a common catalytic triad constituting Cys138, His139, and Glu170. A characteristic hallmark of DJ-1 superfamily members is the redox sensing catalytic cysteine, whose oxidation state determines the physiological function (Wilson, 2011). The emerging evidence on bacteria, humans, and plant DJ-1 members reveals a unique deglycase machinery that relieves the MG and GO glycation adducts on DNA and proteins (Prasad et al., 2022; Richarme et al., 2017; Smith et al., 2022). This repair mechanism is further appreciated in preventing protein aggregation and altering the epigenetic landscape of the chromatin (Sharma, Rao, & Kalivendi, 2019; Zheng et al., 2019). Some *S. cerevisiae* Hsp31 members are reported to possess crucial functions in regulating ROS, mitochondrial dynamics, and chaperone activity (Aslam, Tsai, & Hazbun, 2016; Bankapalli et al., 2020; Miller-Fleming et al., 2014; Tsai et al., 2015). However, the biological relevance of having multiple DJ-1 orthologs in *S. cerevisiae* is poorly understood due to the lack of *in vitro* evidence and difficulties in targeting specific paralog as they are genetically almost identical.

In the current report, we have uncovered a primary role of the Hsp31 paralogs as glyoxalases and protectants of the genome against genotoxic agents. They prevent macromolecular glycation by governing endogenous dicarbonyl levels through scavenging MG and GO. Moreover, the glycation of RNA significantly reduced mRNA translation activity in the absence of yeast DJ-1 members. Additionally, the novel deglycase machinery efficiently repairs glycated DNA and proteins that substantially revert damaged biomolecules. Furthermore, our data highlights carbonyl stress-dependent redistribution of yeast DJ-1 orthologs into mitochondria. Together, the dual role is critical in providing multifaceted protection to cellular and mitochondrial macromolecules during persistent carbonyl stress.

## Results

### Deletion of Hsp31 paralogs aggravates carbonyl toxicity and induces proteome glycation

The DJ-1 superfamily members strongly relate to regulating dicarbonyl stress primarily through scavenging excess intracellular MG and GO (J. Y. Lee et al., 2012; Subedi, Choi, Kim, Min, & Park, 2011). However, despite being conserved, the biological significance of *S. cerevisiae* Hsp31 paralogs is uncharacterized in the presence of carbonyl toxicity. Although the paralogs have very similar amino acid sequences, we utilized the ease of genetic manipulation in yeast to unravel their association with the homeostasis of RCS (*Figure 1-figure supplement 1A*). Hsp31 paralogs were sequentially deleted in the WT background in multiple combinations such as Δ*hsp31* (Δ*31*) to Δ*hsp34 (*Δ*34*), Δ*31*Δ*32* to Δ*31*Δ*34*, Δ*31*Δ*32*Δ*33* (Δ*T),* and Δ*31*Δ*32*Δ*33*Δ*34* (Δ*Q*) which simplifies the comprehensive understanding of the individual genes. As glyoxalase-1 (GLO1) is extensively involved in modulating endogenous RCS levels, Δ*glo1* strain was used for comparative analysis (Gomes et al., 2005). Phenotypic analyses under GO stress revealed a mild growth sensitivity in single deletion strains (*Figure 1A, compare panels glyoxal (GO) with control*), and the viability was further affected in Δ*31*Δ*32,* Δ*31*Δ*33,* and Δ*31*Δ*34.* Interestingly, Δ*T and* Δ*Q* displayed synergistic worsening of growth compared to other strains, inferring their additive overlapping functions during GO toxicity. Subsequently, the effect of MG stress was tested in the deletion background of Hsp31 members. Among the paralogs, Δ*31* alone displayed mild growth sensitivity, as reported earlier (*Figure 1A, compare panels methylglyoxal (MG) with control*) (Bankapalli et al., 2015). On the other hand, additional loss of Hsp32, Hsp33, and Hsp34 in Δ*31* had no cumulative effect on viability.

**Figure 1.**
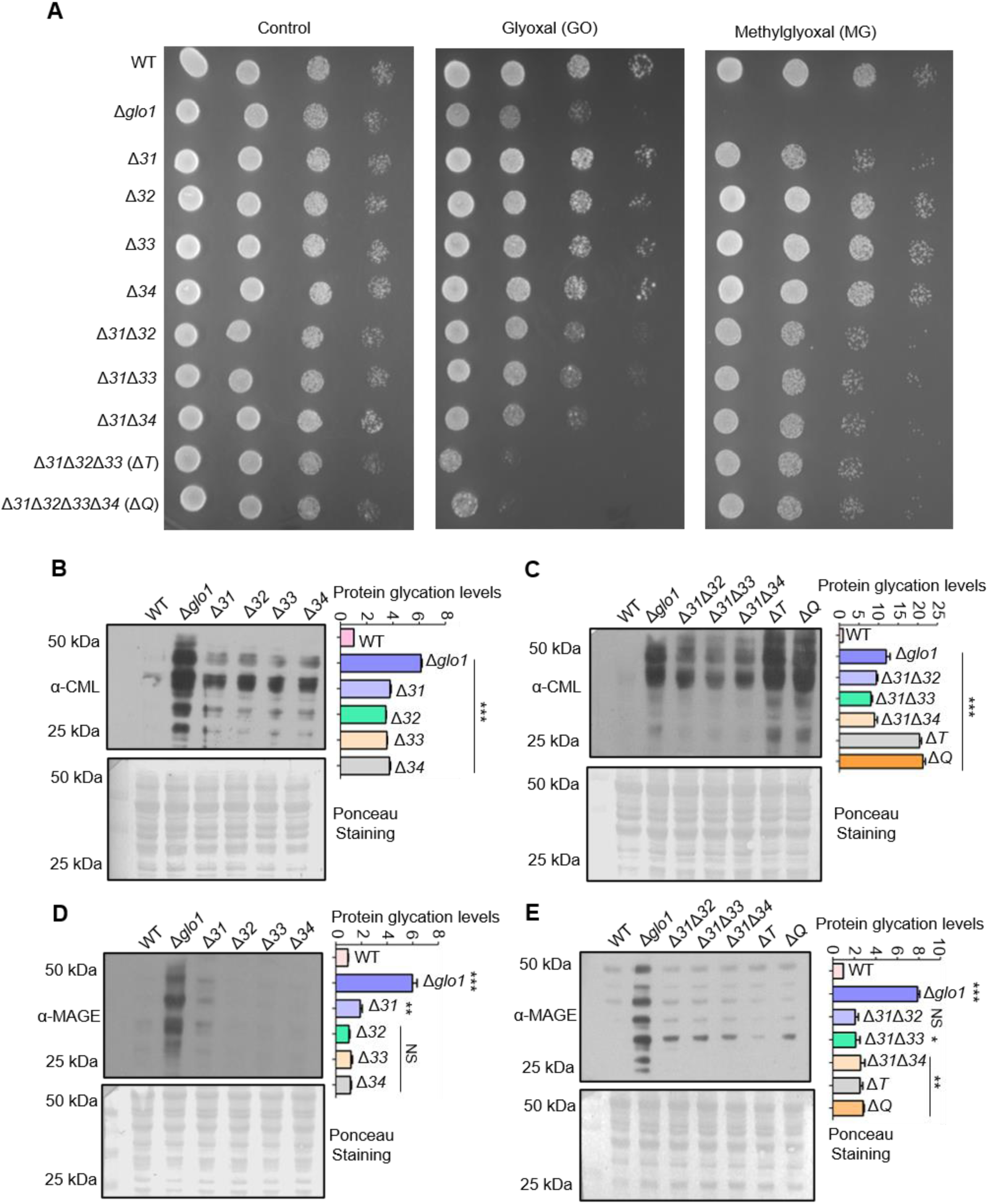
Deletion of Hsp31 paralogs induces protein glycation. (**A**) Yeast phenotypic analysis. Cells were grown until the mid-log phase and harvested, subsequently treated with 10 mM MG before being spotted or spotted on YPD medium plates containing 15 mM GO. The plates were incubated at 30⁰C and imaged at 36 h. (**B, C**) Proteome glycation profile. Yeast strains were treated with 15 mM GO in the YPD culture medium and allowed to grow for 12 h, followed by western analysis with *a*nti-CML antibody. (**D, E**) MAGE detection by western blotting. Cells from the mid-log phase were incubated with 10 mM MG for 12 h, and MAGE levels were estimated using an anti-MAGE antibody. Protein glycation levels were determined by measuring the whole lane intensities by densitometry and plotted with respect to WT. The blots stained with Ponceau S were used as the loading control. One-way ANOVA with Dunnet’s multiple comparisons test was used to determine significance(n=3), *, *p*≤ 0.05; **, *p*≤0.01; ***, *p*≤ 0.001; NS, not significant. **Figure 1 —source data 1.** Source data for *Figure 1B* **Figure 1 — source data 2.** Source data for *Figure 1C* **Figure 1 — source data 3.** Source data for *Figure 1D* **Figure 1 — source data 4.** Source data for *Figure 1E*

To test whether the observed phenotypes are the consequence of impaired carbonyl homeostasis, protein modifications were examined in the deletion background of Hsp31 paralogs. To determine the GO and MG-induced alterations, anti-CML (carboxymethyl-lysine) and anti-MAGE (methylglyoxal AGE) antibodies were utilized to determine the intermediates of AGEs, respectively. The basal physiological glycation levels in (absence of glycation stress), Δ*Q* strain displayed an increment in GO-modified proteins compared to Δ*glo1* strain suggesting Hsp31 paralogs are essential for the RCS detoxification *in vivo* (*Figure 1-figure supplement 1B, C*). However, such basal glycation levels are tolerable to the cells without exhibiting growth sensitivity. Intriguingly, upon subjecting to additional external glycation stress, significant elevation of CML levels was observed in the single mutants compared to WT (*Figure 1B*). At the same time, the deletion of additional members substantially further exacerbated the modifications. It cumulatively enhanced the protein glycation levels to a greater extent in Δ*T* and Δ*Q* than Δ*glo1,* consistent with growth sensitivity (*Figure 1C*). In contrast, the effect of MG-induced glycation was restricted to Δ*31* with a subtle increment of MAGE levels compared to WT, consistent with growth sensitivity in the presence of MG (*Figure 1D and compare growth in the last panels, Fig 1A*). Intriguingly, the collective loss of the members did not exhibit additional MAGE modification in agreement with the growth sensitivity in the presence of MG (*Figure 1E and compare growth in the last panels, Fig 1A*). This preliminary data demonstrate that yeast DJ-1 members act on specific RCS and collectively provide superior tolerance towards GO toxicity than the GLO1. Strikingly, Hsp31 alone imparts resistance for both GO and MG, together preventing the persistent ongoing protein glycation.

### The absence of Hsp31 paralogs enhances nucleic acid glycation and abates mRNA translation efficiency

The nucleic acids are highly susceptible to modifications and are constantly challenged with several metabolic by-products that elicit mutations and chromosomal aberrations (Chatterjee & Walker, 2017). Occasionally, the alterations are irreversible and may not be repaired despite stringent repair systems, culminating to disease conditions. Since the absence of Hsp31 paralogs augments protein glycation, the glycation status of cellular nucleic acids was also assessed. Despite the absence of GO stress, elevation in DNA glycation levels was observed in both Δ*T* and Δ*Q* (*Figure 2-figure supplement 1A, B*). However, in the presence of GO stress, the single deletion strains exhibit further enhancement in the glycation compared to WT (*Figure 2A*). At the same time, the additional loss of Hsp31 paralogs led to a synergistic increment of DNA adducts, most evident in Δ*T* and Δ*Q* strains consistent with the growth sensitivity (*Figure 2B*).

**Figure 2.**
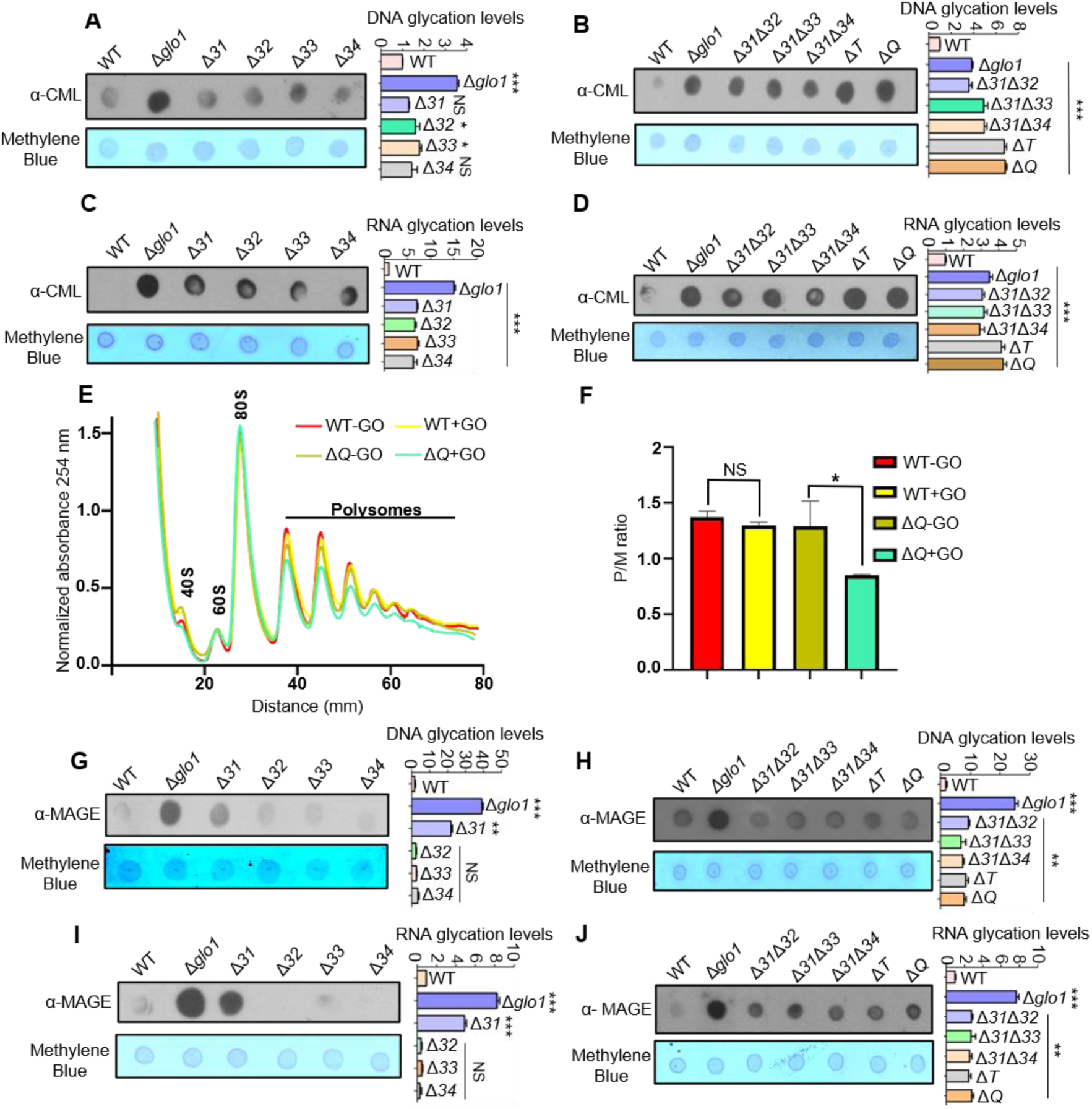
Loss of Hsp31 paralogs aggravates the glycation of DNA and RNA, affecting translational activity. **(A, B)** Immunodetection of DNA using dot-blot assay. 3 µg of total genomic DNA from respective strains treated overnight with 15 mM GO was dotted on nitrocellulose membrane and probed with *a*nti-CML antibody. **(C, D)** RNA glycation profile. Global RNA extracted from cells incubated with 15 mM GO were dotted and analyzed using *a*nti-CML antibody. **(E)** Polysome profiling. Untreated (-GO) and treated (+GO) WT and Δ*Q* cells with GO were subjected to polysome profiling. **(F)** Ratio of polysomes to monosomes (P/M). The area occupied by the polysome and the monosome peak was determined using Origin 8.0 for the respective samples, and the ratios were plotted. WT (+GO) was compared with WT (-GO), and Δ*Q (*-GO) was compared with Δ*Q* (+GO). **(G-J)** Estimation of MAGE-modified nucleic acids. Strains lacking Hsp31 paralogs were treated with 10 mM MG for 12 h, and the genomic DNA and RNA were dotted on the membrane and probed with anti-MAGE antibody. The relative intensity of dots representing the glycation levels was calculated and compared to WT. One-way ANOVA with Dunnet’s multiple comparisons test was used to determine significance (n=3), *, *p*≤ 0.05; **, *p*≤0.01; ***, *p*≤ 0.001; NS, not significant. **Figure 2— source data 1.** Source data for *Figure 2A* **Figure 2— source data 2.** Source data for *Figure 2B* **Figure 2 — source data 3.** Source data for *Figure 2C* **Figure 2— source data 4.** Source data for *Figure 2D* **Figure 2— source data 5.** Source data for *Figure 2E* **Figure 2— source data 6.** Source data for *Figure 2F* **Figure 2— source data 7.** Source data for *Figure 2G* **Figure 2— source data 8.** Source data for *Figure 2H* **Figure 2— source data 9.** Source data for *Figure 2I* **Figure 2— source data 10.** Source data for *Figure 2J*

Besides the alterations on DNA, glycation of RNA was also determined, as its modifications lead to ribosome stalling, translational defects, and inefficient binding with proteins (Mitchell et al., 2018; Zheng, Maksimovic, Upad, & David, 2020). Like DNA, the lack of Hsp31 paralogs led to the high frequency of RNA modifications, which were quantitatively higher than Δ*glo1* (*Figure 2C, D*). Furthermore, the effect of glycation on global mRNA translational activity was evaluated between WT and Δ*Q*. In the absence of GO stress, WT and Δ*Q* showed similar polysome profiles (*Figure 2E*). However, the mRNA translational activity was significantly reduced in Δ*Q* treated with GO, as indicated with low-intensity polysome peaks (*Figure 2E*). Moreover, the ratio of polysome to monosomes also suggested a reduced translation rate in ΔQ compared to WT in the presence of glycation stress (*Figure 2F*).

Next, the role of yeast DJ-1 orthologs in maintaining genome integrity was investigated during MG stress. Unlike Δ*31*, the DNA glycation in Δ*32,* Δ*33*, and Δ*34* was minimally detected by α-MAGE (*Figure 2G*), inferring their lack of participation in MG detoxification. Further, the paralogs were cumulatively deleted to evaluate the possibility of additive effects on glycation levels. Interestingly, the MAGE-modified DNA was restricted to the loss of Hsp31 due to its role in MG homeostasis (*Figure 2H*). Also, deletion of Hsp31 alone led to substantial glycation of RNA when treated with MG (*Figure 2I, J*). In summary, our results highlight that the DJ-1 paralogs in yeast play an essential role in modulating the RCS-mediated glycation of DNA and RNA.

### Yeast DJ-1 orthologs are robust glyoxalases that attenuate the glycation of macromolecules

DJ-1 superfamily members are reported to detoxify MG and GO independently by the same active site with varying kinetics(Bankapalli et al., 2015; J. Y. Lee et al., 2012). Hence, to establish a direct correlation of enhanced glycation with excess intracellular RCS, Hsp31 paralogs were purified using affinity chromatography and their glyoxalase activity was determined *in vitro.* Remarkably, our *in vitro* experiments suggested that Hsp31, Hsp32, Hsp33, and Hsp34 possess a robust glyoxalase activity compared to BSA, which served as negative control (*Figure 3A*). Furthermore, the kinetic parameters indicate that paralogs have relative catalytic efficiencies. Notably, the paralogs exhibited 4-fold higher *k_cat_* for glyoxalase activity than hDJ-1 (J. Y. Lee et al., 2012) (*Figure 3B* and *Figure 3-figure supplement 1A*). To further appreciate their *in vivo* biological response during GO stress, Hsp31 members were overexpressed within Δ*Q* strain and observed a substantial rescue in the growth similar to WT (*Figure 3C*). The expression of the proteins was normalized to equal amounts (*Figure 3-figure supplement 1B)*.

**Figure 3.**
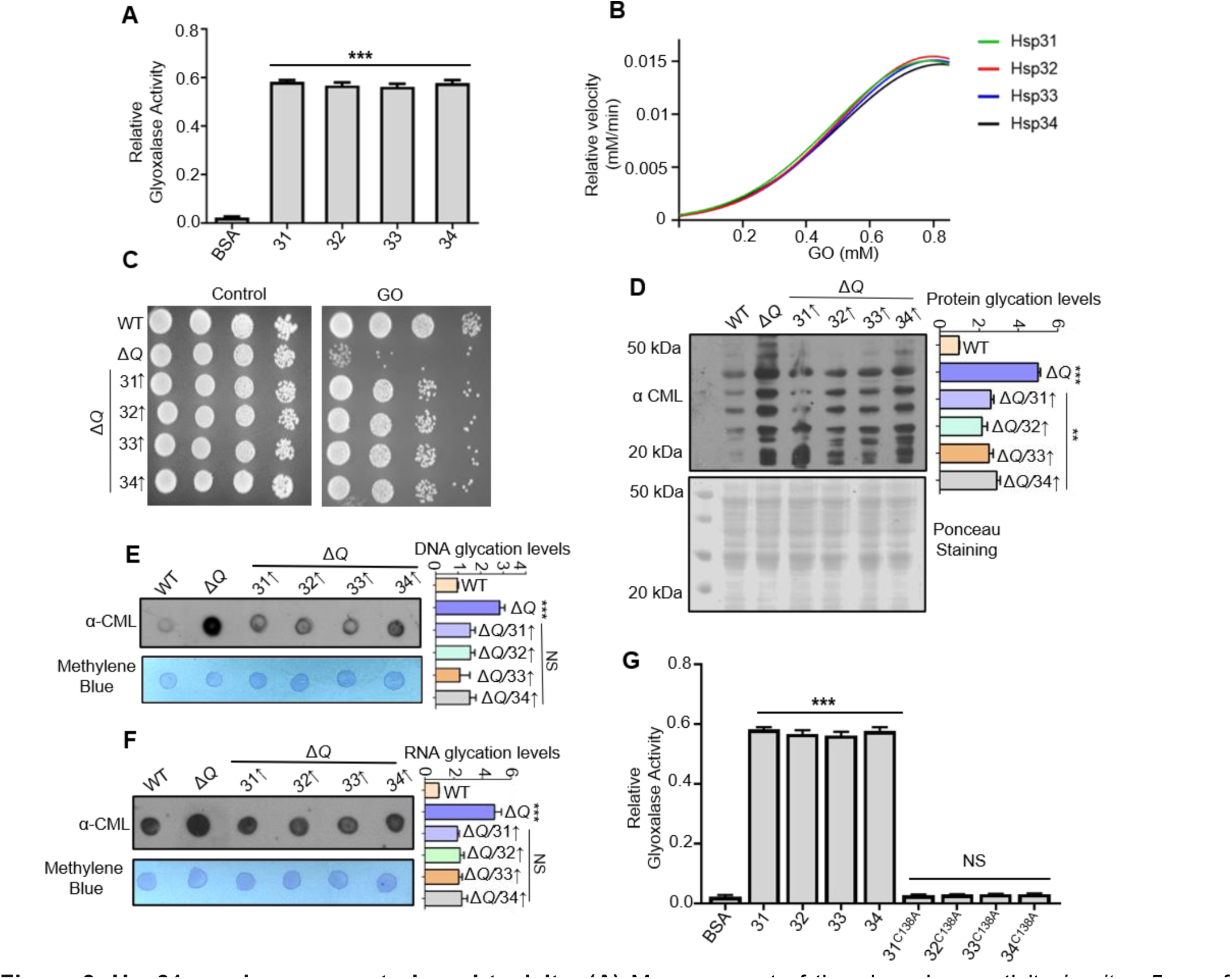
Hsp31 paralogs prevent glyoxal toxicity. **(A)** Measurement of the glyoxalase activity *in vitro*. 5 µg of purified Hsp31 paralogs were incubated with 0.5 mM GO, and the absorbance was monitored at 570 nm. **(B)** Estimating the kinetic parameters of the enzyme. The kinetic efficiency was determined by calculating their reaction velocities against multiple GO concentrations. **(C)** Growth phenotypic analysis. WT and Δ*Q* overexpressing Hsp31 paralogs were grown until the mid-log phase, spotted on SD Leu-media plates containing 15 mM GO, and incubated at 30⁰C. **(D)** Yeast DJ-1 homologs alleviate proteome glycation. Respective strains were grown in SD Leu-culture tubes and treated with 15 mM GO for 12 h, and the whole cell lysates were processed and probed for CML modifications. **(E, F)** Analysis of nucleic acid modifications by dot-blot assay. Genomic DNA and global RNA extracted from cells treated with 15 mM GO were dotted on the membrane and immunoreacted with *a*nti-CML-antibody. **(G)** C138 amino acid residue is essential for glyoxalase activity. All the proteins were purified and incubated (5 µg) with 0.5 mM GO; subsequently, the absorbance was monitored at 570 nm. Significance was determined by Dunnett’s multiple comparison test with WT or BSA (enzyme activity). Glycation levels represent the relative intensities of the dots and lanes compared with WT. One-way ANOVA with Dunnet’s multiple comparisons test was used to determine significance (n=3), **, *p*≤0.01; ***, *p*≤ 0.001; NS, not significant. **Figure 3— source data 1.** Source data for *Figure 3A* **Figure 3— source data 2.** Source data for *Figure 3B* **Figure 3— source data 3.** Source data for *Figure 3D* **Figure 3— source data 4.** Source data for *Figure 3E* **Figure 3— source data 5.** Source data for *Figure 3F* **Figure 3— source data 6.** Source data for *Figure 3G*

To further corroborate the above findings, Δ*Q* strain overexpressing Hsp31 paralogs was subjected to GO stress and probed for CML levels. In agreement with our data, the modifications on proteins and nucleic acids were substantially suppressed in the presence of yeast DJ-1 glyoxalase machinery (*Figure 3D-F*). Next, the role of nucleophile cysteine amino acid (Cys-138) in the glyoxalase activity was tested, as its mutations predominantly impair the functions of DJ-1 members (Bankapalli et al., 2015; J. Y. Lee et al., 2012; Zheng et al., 2019). Therefore, cysteine 138 was replaced with alanine (C138A), and the proteins were subjected to the glyoxalase activity assay. Upon *in vitro* analysis, all the mutant proteins had compromised activity suggesting the critical requirement of cysteine in mitigating intracellular carbonyl levels (*Figure 3G*). Furthermore, we observed that exposure of cells to excess GO concomitantly enhanced the steady-state levels of Hsp31 paralogs approximately to three-fold (*Figure 3-figure supplement 1C, D*). These results indicate a direct role of yeast DJ-1 orthologs in modulating CML levels through regulating endogenous glyoxal.

### Hsp31 alone specifically prevents the accretion of methylglyoxal-derived AGEs

MG is the most reactive dicarbonyl with the highest potency for glycating vital macromolecules thereby causing organellar damage due to membrane permeability (Allaman et al., 2015).To overcome the MG toxicity, the glyoxalase system (GLO1 and GLO2) inevitably exploits the GSH pool for detoxification and thus alters the cellular redox balance (de Bari et al., 2020). On the other hand, DJ-1 members are functionally independent and do not utilize GSH as a cofactor for detoxification. To determine the methylglyoxalase activity of Hsp31 paralogs *in vitro*, we purified proteins recombinantly and subjected them to activity analysis. Strikingly, Hsp31 alone exhibited robust methyglyoxalase activity compared to other paralogs (*Figure 4A*). This further confirms the specific enrichment of MAGE modification reported earlier Δ*31* strain (*Figure 1D, 2G, and 2H*). On the contrary, Hsp32, Hsp33, and Hsp34 demonstrated significantly lower methyglyoxalase activity in agreement with the lack of MAGE modifications (*Figure 1D, 2G*, and *2H*).

**Figure 4.**
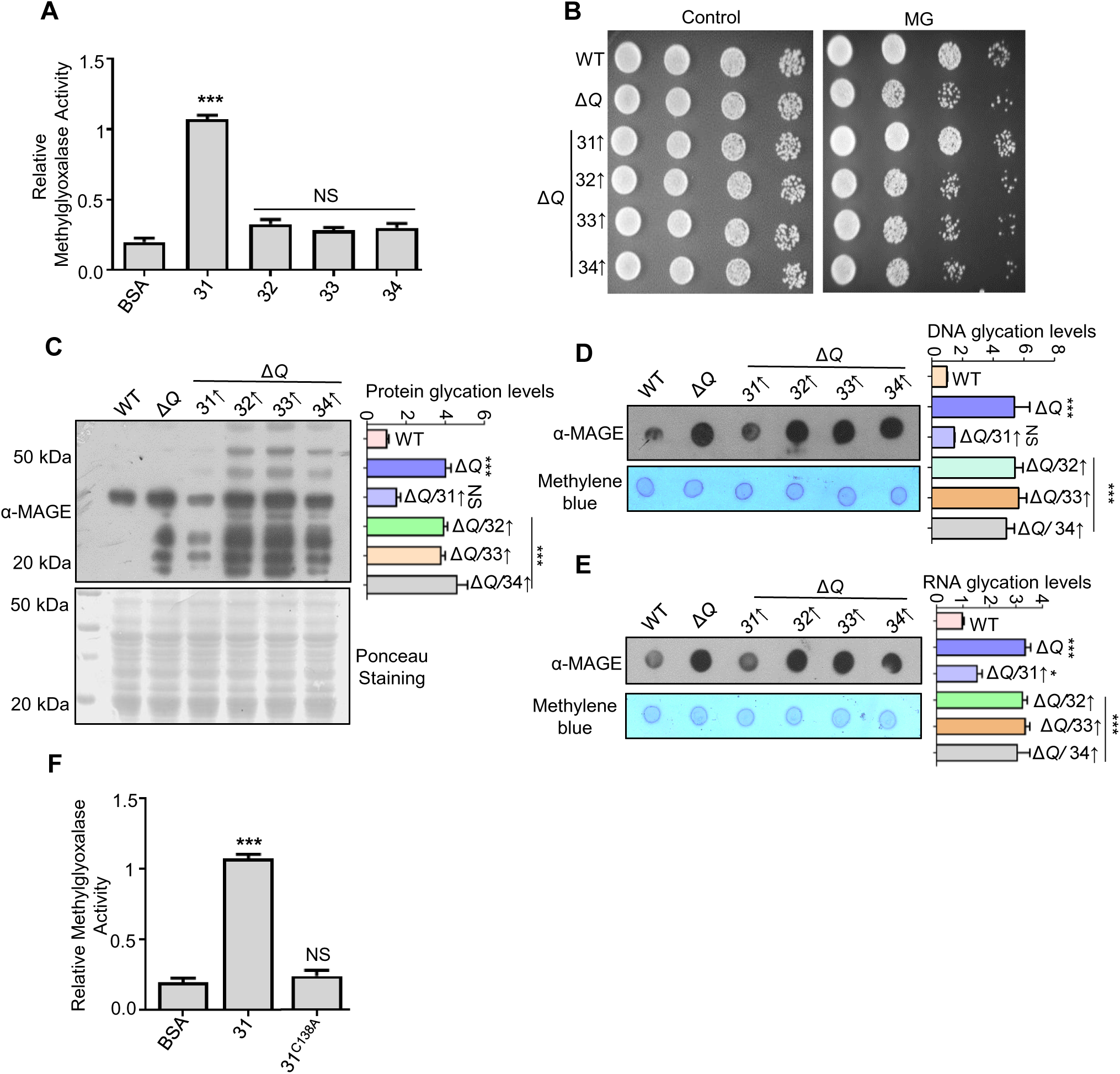
Hsp31 is a potent scavenger of methylglyoxal. **(A)** *In vitro* methylglyoxalase activity. Purified Hsp31 members 5 µg each were incubated with 0.5 mM MG, and the absorbance at 530 nm was monitored. **(B)** Spot assay. Respective strains grown till the mid-log phase were harvested and treated with 10 mM MG for 5 h before spotting on SD Leu-plates. Images were captured at 36 h. **(C-E)** Hsp31 reduces MG-derived AGE modifications. Individual strains were treated with 10 mM MG in the culture tubes for 12 h, and the macromolecule glycation was analyzed using an anti-MAGE antibody. **(F)** C138 amino acid residue is critical for methylglyoxalase activity. 5 µg of Hsp31-WT and Hsp31^C138A^ mutant were incubated with MG, and the activity was examined at 530 nm. Significance was determined by Dunnett’s multiple comparison test with WT or BSA (enzyme activity). Glycation levels represent the relative intensities of the dots and lanes compared with WT. One-way ANOVA with Dunnett’s multiple comparisons test was used to determine significance (n=3), *, *p*≤ 0.05; ***, *p*≤ 0.001; NS, not significant. **Figure 4—source data 1.** Source data for *Figure 4A* **Figure 4—source data 2.** Source data for *Figure 4C* **Figure 4—source data 3.** Source data for *Figure 4D* **Figure 4—source data 4.** Source data for *Figure 4E* **Figure 4—source data 5.** Source data for *Figure 4F*

Remarkably, the overexpression of Hsp31 in Δ*Q* strain facilitated improved resistance towards MG treatment than its paralogs due to its greater methyglyoxalase activity (*Figure 4B*). Further, examining the proteome glycation by α-MAGE also indicated that Hsp31 markedly reduced MG glycated proteins compared to its paralogs (*Figure 4C*). To further substantiate the implications of methylglyoxalase activity of Hsp31 paralogs, modification of nucleic acids from the respective strains was investigated. In agreement with high intrinsic methylglyoxalase activity, Hsp31 alone was able to scavenge MG-associated glycation of nuclear DNA and global RNA (*Figure 4D, E*). Besides, the role of a catalytic cysteine (C138) residue in methylglyoxalase activity was evaluated due to its pivotal role in abating GO toxicity. The mutation of C138 to alanine in Hsp31 completely abolished the enzymatic activity (*Figure 4F*), which further emphasizes the prominence of a conserved cysteine in the catalytic triad of DJ-1 superfamily members. Together, our data suggest that Hsp31 provides glycation defense against both MG and GO, unlike Hsp32, Hsp33, and Hsp34.

### Hsp31 paralogs repair glyoxal glycated DNA and proteins, suppress genetic mutations, and combat genotoxic stress

Glycation of macromolecules in *S. cerevisiae* is a rapid and spontaneous event with specific targets allowing them to tune their physiological functions (Gomes et al., 2005). However, glycation is also an irreversible process that may erroneously alter the features of biomolecules in the absence of a dedicated restoration pathway. Emerging studies have reported hDJ-1 as deglycase (EC 3.5.1.124) that relieve advanced glycation adducts from MG and GO-modified nucleotides and proteins (Richarme et al., 2017; Richarme et al., 2015). This observation unravelled a gateway to regulating the glycation status of essential components within cells across species.

We utilized the modified protocols to test whether the yeast Hsp31 paralogs possess deglycase activity (Prasad et al., 2022; Richarme et al., 2017). Briefly, deoxyribonucleotide triphosphate (dNTPs) were pre-treated without or with GO to completely glycate the substrates. The glycation mixture was further diluted to 6-fold to suppress the effect of residual GO on the activity of Hsp31 paralogs. Later, the glycated substrates were incubated for 3 h with purified Hsp31 paralogs and BSA as a control to test deglycase activity (*Experimental scheme; Figure 5A*). The deglycase activity was ascertained by amplifying the *S. cerevisiae sod1* gene generated by Phusion high fidelity DNA polymerase that incorporates repaired dNTPs during PCR reaction. Unlike GO and BSA controls, incubation with Hsp31 paralogs efficiently reverted the glycation adducts of dNTPs, noted by robust amplification of the PCR product (*Figure 5B*). As an additional substrate of DNA, the forward and reverse oligonucleotide primers of the *sod1* gene were glycated in the presence or absence of GO. Besides dNTPs, yeast DJ-1 orthologs prominently deglycated the primers, allowing the reaction to yield an intense PCR product in contrast to GO and BSA controls (*Figure 5-figure supplement 1A*).

**Figure 5.**
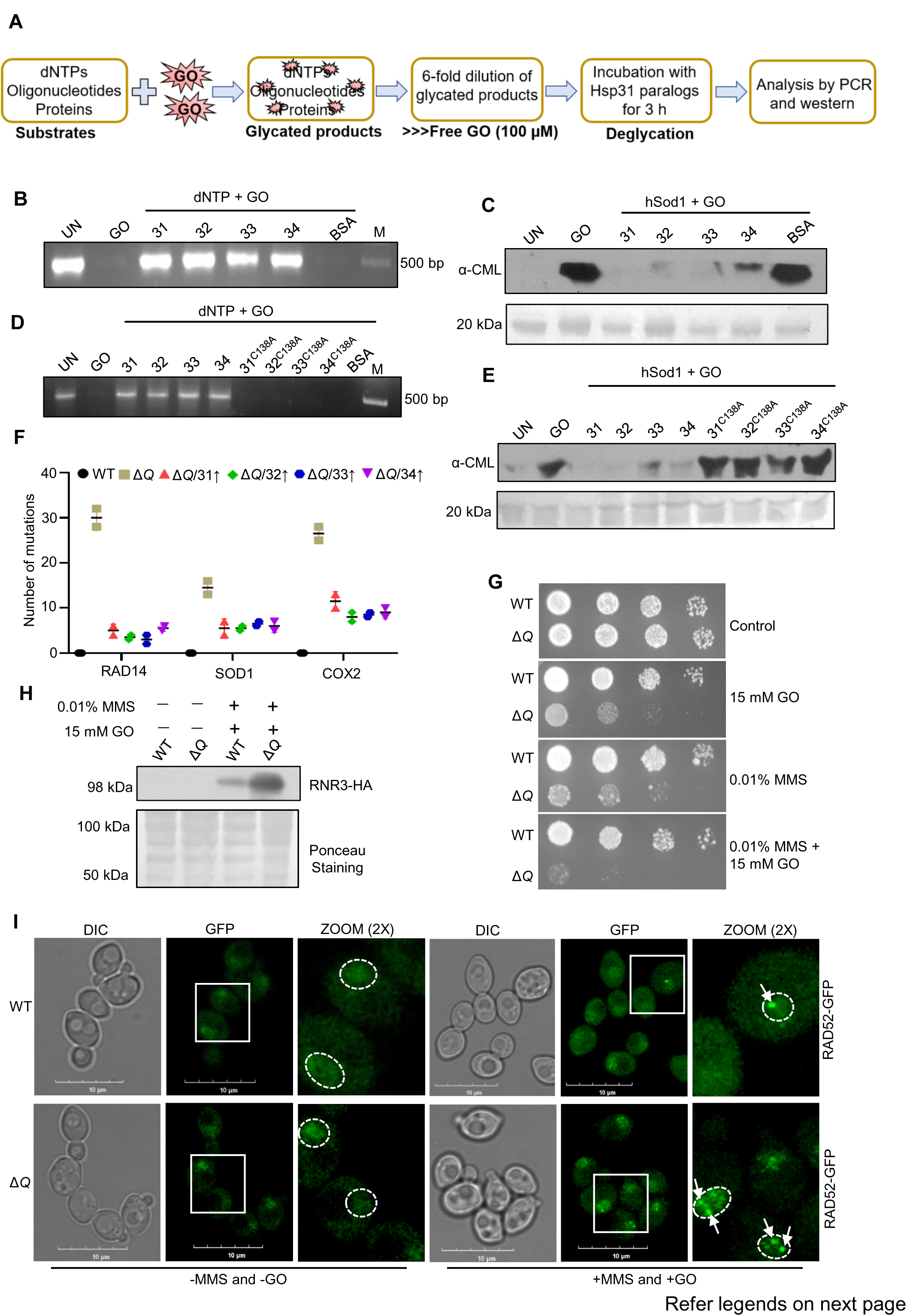
*In vitro* deglycation of DNA and proteins, and genotoxic sensitivity in the absence of yeast DJ-1 orthologs. **(A)** Schematic representation of the experimental procedure followed for DNA and protein deglycation reactions. **(B)** Hsp31 paralogs deglycate DNA. 500 µM dNTPs were incubated without (UN) or with 2 mM GO for 2 h, followed by 3 h incubation with 5 µg Hsp31 paralogs. The samples were examined for deglycation through PCR. **(C)** Yeast DJ-1 members repair glycated proteins. 2 µg of purified protein hSod1 was treated without (UN) or with 2 mM GO for 2 h, and the reactions were further incubated for 3 h with Hsp31 paralogs. An anti-CML antibody was used to determine glycation levels. **(D-E)** C138 amino acid residue is crucial for deglycase activity. Glycated macromolecules were individually supplemented with 5 µg of Hsp31 paralogs (WT), or mutant paralogs (C138A) and subjected to PCR or western analysis to determine the glycation status of DNA and proteins, respectively. BSA was used as a negative control in all experiments. 10 kb DNA ladder was used as a marker (M) for DNA gels, and the Ponceau S stain indicates the equal loading of protein samples. **(F)** Hsp31 paralogs attenuate genetic mutations. Individual genes were PCR amplified, and Sanger sequenced from the isolated genome of strains treated with 15 mM GO. The number of genetic mutations was calculated and plotted on GraphPad prism 5.0. **(G)** Growth phenotypic analysis. Cells grown until the mid-log phase were spotted on plates containing 0.01% MMS or 15 mM GO or 0.01% MMS and 15 mM GO. The plates were incubated at 30⁰C and imaged at 36 h. **(H)** Western analysis of RNR3 levels. WT and Δ*Q* grown till the mid-log phase were supplemented with 0.01% MMS, and 15 mM GO in the culture media and further incubated for 3 h. Subsequently, RNR3 levels were probed using an anti-HA antibody. **(I)** RAD52 foci formation. Cells were grown until the mid-exponential phase and were treated without (-) or with 0.03% MMS and 15 mM GO for 1 h. Subsequently, cells were imaged using a confocal microscope (Olympus FV3000); representative images have a 10 µm scale. **Figure 5—source data 1.** Source data for *Figure 5B* **Figure 5—source data 2.** Source data for *Figure 5C* **Figure 5—source data 3.** Source data for *Figure 5D* **Figure 5—source data 4.** Source data for *Figure 5E* **Figure 5—source data 5.** Source data for *Figure 5F*

The chronic glycation of proteins induces multifactorial changes that potentially trigger aggregation, cross-linking and cytotoxicity. Therefore, it is critical to study whether Hsp31 paralogs confer the glycation repair of proteins. To determine the deglycase property, human Sod1 and Lysozyme were pre-treated without or with GO separately for 2 h to form the glycation adducts on proteins. The reaction mixture was diluted 6-fold and incubated for 3 h with purified Hsp31 members or BSA (negative control) (*Experimental scheme; Figure 5A*). The samples were separated on an SDS-PAGE and subjected to immunodetection by anti-CML antibody to ascertain the degree of glycation. All the Hsp31 members exhibited robust deglycase activity, as the paralogs extensively reverted the glycation damage of proteins, unlike GO and BSA controls (*Figure 5C* and *Figure 5-figure supplement 1B*). On the other hand, the active site mutation (C138A) in paralogs was tested to verify deglycase enzyme activity further. The active site mutants failed to relieve the glycation adducts on DNA and proteins, thus establishing the specificity of the deglycase action of Hsp31 paralogs (*Figure 5D, E* and *Figure 5-figure supplement 1C, D*).

GO is known to induce random mutations that severely impair genome integrity. We scored the glycation-induced conversions at the DNA level to investigate the impact of glyoxalase and deglycase activity of Hsp31 paralogs on genome protection. Strains lacking Hsp31 paralogs were subjected to GO treatment, and mutation frequency was determined using amplification of specific genes such as mitochondrial-encoded (*cox2*) and nuclear-encoded (*sod1* and *rad14*), which have vital *in vivo* functions. Owing to the complete lack of Hsp31 paralogs, Δ*Q* strain exhibited enhanced mutation frequency as determined by sequencing of *cox2*, *sod1*, and *rad14* genes (*Figure 5F)*. Intriguingly, due to robust intrinsic glyoxalase and deglycase activity associated with the paralogs, the overexpression of Hsp31, Hsp32, Hsp33, and Hsp34 significantly attenuated the mutation frequency of all the genes tested, further highlighting their physiological role in the maintenance of genome integrity under the GO toxicity (*Figure 5F and Figure 5-figure supplement 1E*).

To establish their role further in genome protection, WT and Δ*Q* strains were exposed to well-known genotoxic agents such as methyl methanesulfonate (MMS) and hydroxyurea (HU). The Δ*Q* strain showed severe growth sensitivity towards MMS and HU treatment compared to WT (*Figure 5-figure supplement 1F*). Furthermore, treatment with GO and genotoxins combined showed a synergistic increment in the growth sensitivity of Δ*Q* compared to their individual treatments (*Figure 5G*). These genotoxins induce DNA lesions and strand breaks, which could be monitored by the expression of DNA damage response genes. Therefore, we determined the expression of RNR3 (subunit of RiboNucleotide Reductase), a DNA damage marker protein that specifically upregulates during genotoxic stress (Fu, Pastushok, & Xiao, 2008). Intriguingly, Δ*Q* strain exhibited a notable increase in the expression of RNR3 in contrast to WT cells treated with MMS and GO (*Figure 5H*). Besides RNR3, multiple DNA repair proteins, including RAD52, assemble at the strand break regions. Hence, the formation of RAD52-GFP foci indicates the extent of DNA damage in cells (Conde & San-Segundo, 2008). In support of our findings, ΔQ strain displayed a more significant number of RAD52 foci, with cells showing more than one focus per cell compared to WT upon treatment with MMS and GO (*Figure 5I*). In summary, our findings indicate that the Hsp31 members collectively provide genome protection against various genotoxic stresses and participate in an antiglycation pathway that globally detects and restores the GO-modified biomolecules.

### Hsp31 alone explicitly reverts methylglyoxal glycated macromolecules and attenuates mutation frequency

Balancing the physiological amount of dicarbonyls is a major biological challenge as they have a short half-life and quickly react with nearby biomolecules. Besides glyoxalase-I and II systems, very few enzymes have been identified capable of detoxifying MG (Vander Jagt & Hunsaker, 2003). About 90-99% of cellular MG is bound to macromolecules, which may be reversible by the action of deglycases (Allaman et al., 2015). We utilized a modified protocol to test whether the paralogs attenuate MG-mediated modification on DNA and proteins (Richarme et al., 2017). As illustrated in schematic *Figure 6A*, the dNTPs or DNA oligonucleotide primers substrates were treated in the absence or presence of MG, followed by 6-fold dilution of the reaction mixture to reduce the residual effects of MG on the enzymatic activity of Hsp31 paralogs. The glycated mixture was incubated with purified proteins, and deglycase activity was ascertained by PCR of *sod1* gene using Phusion high fidelity DNA polymerase. In line with our findings, Hsp31 alone was able to suppress the MG-mediated glycation through its deglycase activity, and robust amplification of PCR product was observed (*Figure 6B, C*). This is consistent with our previous results where Δ*hsp31* strain displayed MG-mediated growth sensitivity, unlike, Hsp32, Hsp33, and Hsp34, which specifically act on GO glycated macromolecules.

**Figure 6.**
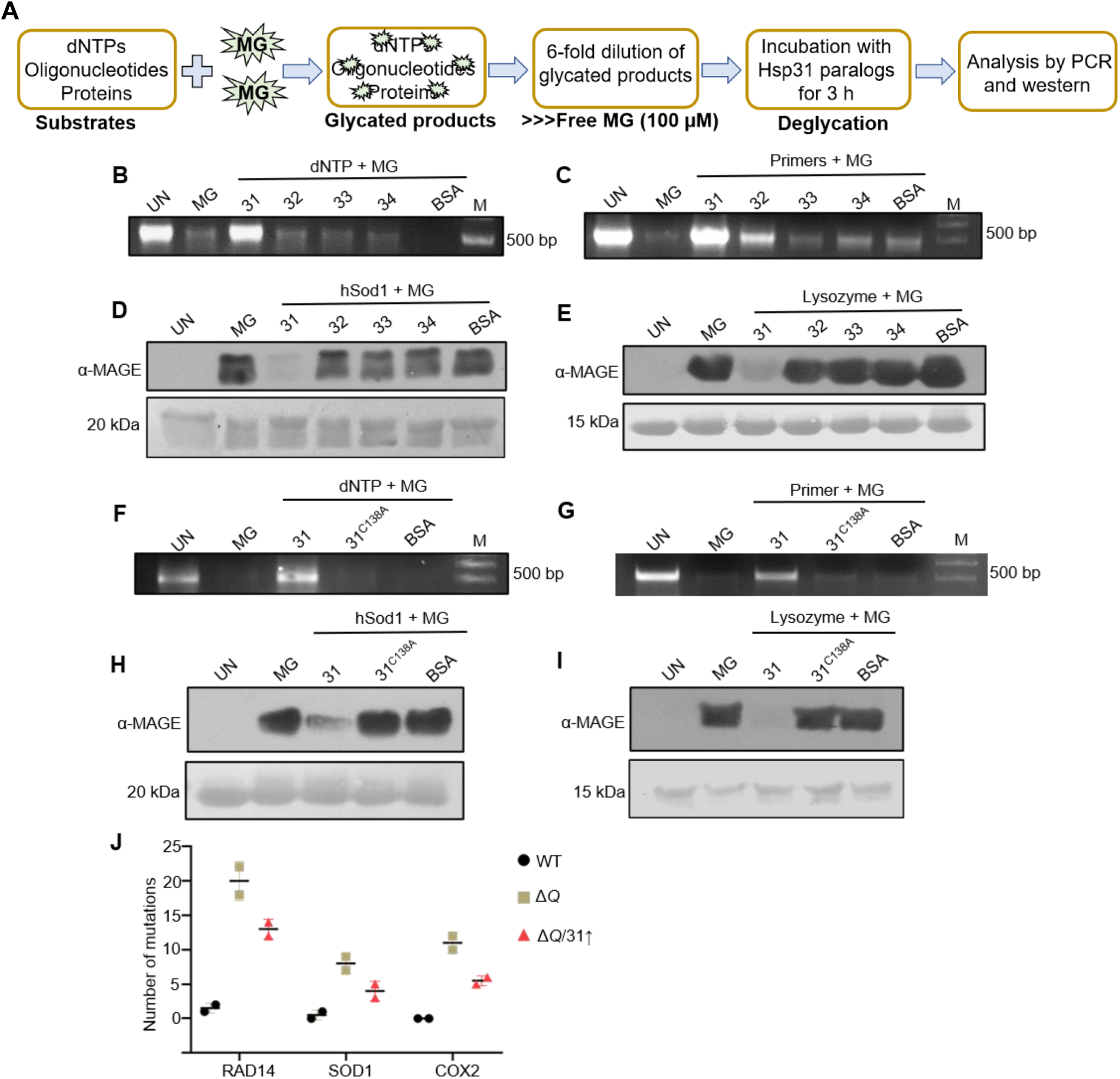
Hsp31 repairs MG-derived AGE modifications on DNA and proteins. **(A)** Schematic representation of the experimental procedure used for DNA and protein deglycation reactions. **(B, C)** DNA deglycation by Hsp31. 500 µM dNTPs or 2 mM primes were incubated without (UN) or with 2 mM MG for 2 h. Subsequently, 5 µg Hsp31 paralogs were added and incubated for 3 h. The samples were subjected for PCR analysis. **(D, E)** Hsp31 reverts MG modification on proteins. hSod1 and Lysozyme were glycated for 2 h with 2 mM MG, followed by incubation with Hsp31 paralogs. The glycation status was determined through western analysis against anti-MAGE antibody. **(F-I)** Hsp31^C138A^ mutant lacks deglycase activity. Macromolecules were treated with 2 mM MG for 2 h, and later incubated with Hsp31-WT and Hsp31^C138A^ mutant (2 µg each). 10 kb DNA ladder was used as a marker (M) for DNA gels, and the Ponceau S stain indicates the equal loading of protein samples. **(J)** The dual role of Hsp31 reduces MG-induced DNA mutations. Respective genes were PCR amplified, and Sanger sequenced from strains treated with 10 mM MG. The number of genetic mutations was plotted on GraphPad prism 5.0. BSA was used as a negative control in all experiments. **Figure 6—source data 1.** Source data for *Figure 6B* **Figure 6—source data 2.** Source data for *Figure 6C* **Figure 6—source data 3.** Source data for *Figure 6D* **Figure 6—source data 4.** Source data for *Figure 6E* **Figure 6—source data 5.** Source data for *Figure 6* **Figure 6—source data 6.** Source data for *Figure 6G* **Figure 6—source data 7.** Source data for *Figure 6H* **Figure 6—source data 8.** Source data for *Figure 6I* **Figure 6—source data 9.** Source data for *Figure 6J*

To delineate the MG-associated protein deglycation, hSod1 and Lysozyme were treated without or with MG and probed with α-MAGE antibody. In the presence of Hsp31, the glycation adducts on the proteins were significantly reverted (*Figure 6D, E*). On the other hand, the Hsp32, Hsp33, and Hsp34 failed to revert the glycation of proteins *in vitro*, similar to DNA substrates (*Figure 6D, E*). Moreover, the Hsp31 active site mutant (C138A) failed to repress the MG adducts on DNA and proteins, underlining its significance in the catalytic reaction (*Figure 6F-I*). We also determined if the dual role of Hsp31 could provide genome protection by scoring the MG-induced genetic mutations. Strikingly, the Δ*Q* strain displayed an augmented mutation rate, which was notably abated by the expression of Hsp31 during MG toxicity as indicated by amplification and sequencing of selected yeast-specific genes *(Figure 6J and Figure 6-figure supplement 1A*). Thus, our data establish a critical function of Hsp31 in suppressing the MG-mediated aberrant glycation of macromolecules in *S. cerevisiae*.

### Dicarbonyl stress induces translocation of Hsp31 paralogs into mitochondria for organelle protection

DJ-1 superfamily members display dynamic subcellular localization into various organelles, such as the nucleus and mitochondria, during the stress response (Bankapalli et al., 2015; Conde & San-Segundo, 2008; Fu et al., 2008). However, the significance of the redistribution is still unclear. Since Hsp31 paralogs attenuate mutations of mitochondrial encoded *cox2* gene, we investigated whether Hsp31 paralogs localize to mitochondria during GO toxicity. To determine, the Hsp31 paralogs were genomically fused with GFP, and the mitochondria were decorated by targeting mCherry tagged with mitochondrial targeting sequence (MTS). Under normal physiological conditions, the paralogs were predominantly localized in the cytosol (*Figure 7A-D*, *-GO untreated panels*). Strikingly, upon treatment of cells with GO, all the Hsp31 paralogs translocate into mitochondria, as highlighted by the merged images of GFP and mCherry (*Figure 7A-D*, +*GO stress panels*). Similarly, our microscopic analysis suggests efficient mitochondrial redistribution of Hsp31 during MG stress (*Figure 7-figure supplement 1A*).

**Figure 7.**
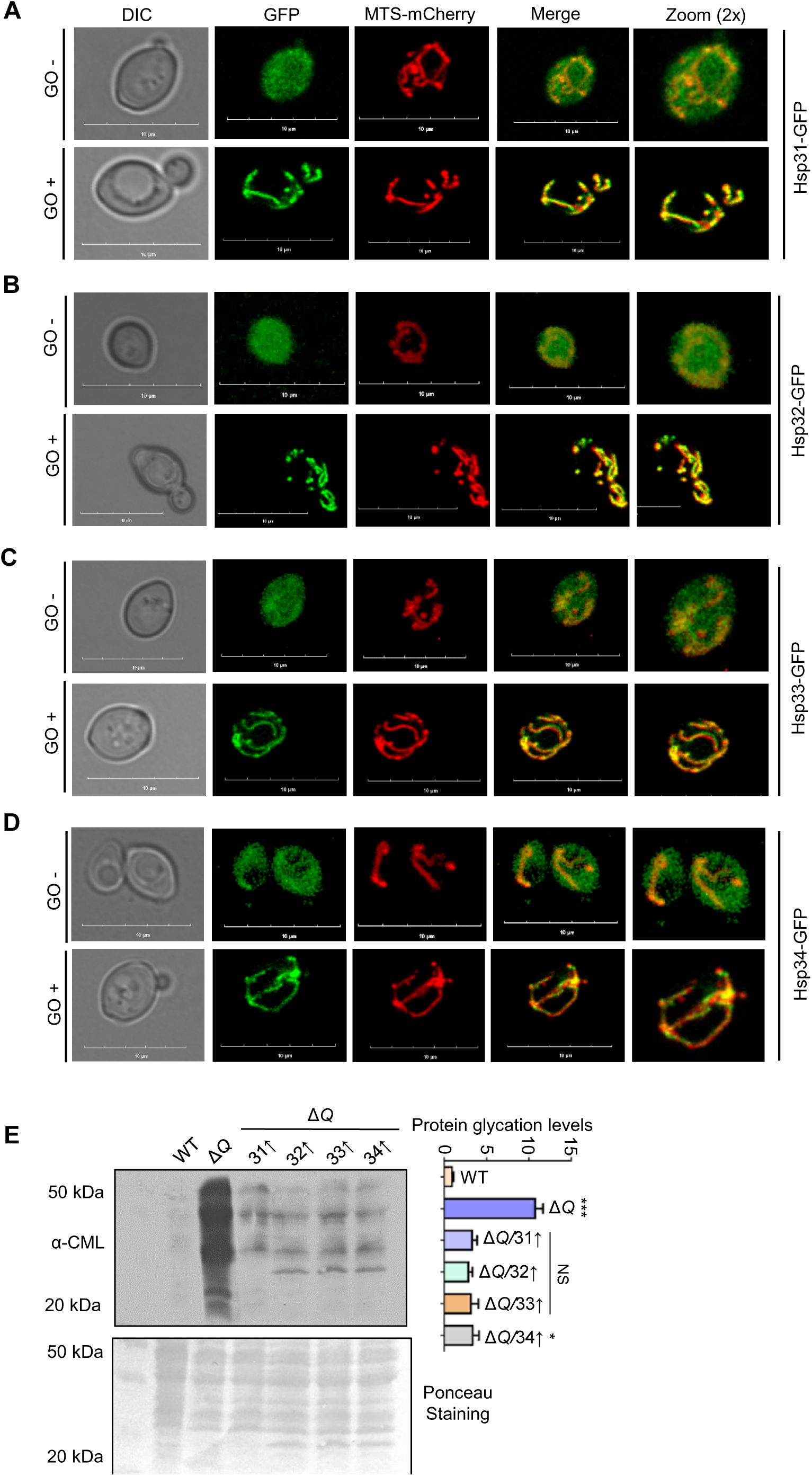
Glyoxal induced translocation of yeast DJ-1 orthologs into mitochondria. **(A-D)** Mitochondrial translocation of Hsp31 paralogs. WT strain expressing MTS-mCherry (decorates mitochondria) and C-terminus GFP tagged Hsp31 paralogs were treated with either buffer (-GO) or 15 mM GO (+GO) for 3 h. Consequently, images were captured in a confocal microscope (Olympus FV3000), and images were represented on a 10 µm scale. **(E)** Mitochondrial protein glycation levels. WT and Δ*Q* overexpressing Hsp31 class of proteins were treated with 15 mM GO stress, followed by isolation of mitochondria and western analysis to determine glyoxal modifications using an anti-CML antibody. The intensity of each lane was quantitated densitometrically and plotted in the graph. One-way ANOVA with Dunnett’s multiple comparisons test was used to determine significance (n=3), *, *p*≤ 0.05; ***, *p*≤ 0.001; NS, not significant. **Figure 7—source data 1.** Source data for *Figure 7E*

To understand the relevance of GO-dependent mitochondrial translocation of Hsp31 paralogs, mitochondria were isolated from cells treated with GO to analyze the altered glycation of mitochondrial proteins. Intriguingly, a significant portion of the mitochondrial proteome was glycated in Δ*Q* strain, and the glycation intensity of proteins was substantially suppressed in the presence of Hsp31 paralogs (*Figure 7E*). In conclusion, our findings indicate that the dual role of Hsp31 paralogs provides robust cellular and organelle protection during dicarbonyl stress.

## Discussion

### Glyoxalase defence mechanism of the Hsp31 class of proteins is more robust than GLO-1 system

The present study highlights robust glyoxalase activities of yeast Hsp31 mini-family proteins essential to combat aberrant intracellular levels of RCS. Our *in vitro* experimental results firmly establish the specificity of Hsp31 paralogs towards GO substrates, while Hsp31 alone detoxifies both MG and GO. Although the paralogs share significant sequence similarities, the Hsp33 possesses an active site different than Hsp31, thus affecting its overall substrate specificity. Due to the 99.5% sequence identity between Hsp33, Hsp32, and Hsp34, their catalytic properties and substrates could be very similar. Such preferential scavenging of carbonyls was also observed in bacteria and plant DJ-1 members where YajL, ElbB, *At*DJ-1A, and *At*DJ-1B are GO specific, and some orthologs detoxify both substrates (Kwon et al., 2013; C. Lee et al., 2016). Although MG and GO are detoxified through distinct pathways, certain NADPH-dependent enzymes like aldehyde dehydrogenase and aldose reductase scavenge both the carbonyls, with their activities stringently governed by cellular redox status (Vander Jagt & Hunsaker, 2003).

In contrast to MG catabolism, antioxidant such as GSH inefficiently reacts with GO making it a poor substrate for the GLO1 glyoxalase system (Yang et al., 2011). Hence, *S. cerevisiae* has bonafide multiple DJ-1 orthologs, and their steady-state levels are rapidly upregulated in response to GO allowing robust carbonyl homeostasis. Our results highlight that deletion of individual paralogs shows moderate sensitivity for GO. However, the collective loss of yeast DJ-1 orthologs (Δ*Q*) renders severe growth deficiency more than Δ*glo1*. At the same time, their absence promotes severe accumulation of toxic carbonyls leading to a higher fold macromolecular glycation than Δ*glo1*. Besides scavenging RCS, Hsp31 is also reported to provide partial tolerance toward acetic acid stress, also a deleterious glycolytic intermediate (Ansari, Chaudhary, & Dash, 2018; Natkańska, Skoneczna, Sieńko, & Skoneczny, 2017). These findings suggest that Hsp31 members are paramount in regulating a broad group of toxic metabolites compared to the GLO system (1 and 2), which is restricted to specific carbonyls. Moreover, under oxidative damage, the activity of GLO system (1 and 2) is impaired due to extensive utilization of the limiting pool of GSH. In contrast, the GSH-independent Hsp31 paralogs aid in restoring redox balance and simultaneously provide robust RCS homeostasis, hence, indispensable for cellular viability under carbonyl stress.

### Global surveillance of nucleic acid glycation by yeast DJ-1 orthologs regulates the genome integrity

Despite several harmful effects, the physiological amounts of glycation are critical for post-translation modification, required for signaling in numerous pathways (Akhand et al., 2001; Rodrigues et al., 2020). Both MG and GO are known to glycate histone proteins. Hitherto, 28 site-specific modifications on histones have been identified as targets of MG (Ansari et al., 2018; Galligan et al., 2018; Ray et al., 2022). Such non-enzymatic covalent epigenetic modifications regulate stress adaptations by altering chromatin architecture and gene expression. We highlight that Hsp31 paralogs are essential repair enzymes that regulate the degree of nucleic acid glycation by indirectly detoxifying the carbonyls or directly removing adducts by deglycation. This corroborates well with the idea that Hsp31 paralogs preserve genome integrity by lowering deleterious mutations as a counter-response (*Figure 8*). Also, the absence of paralogs elicits enhanced sensitivity to DNA damage stress when exposed to multiple genotoxins, as indicated by enriched expression of RNR3 and RAD52 foci formation, suggesting their strong association with genome maintenance. At the molecular level, the antiglycation property of the paralogs could also prevent the replication stress by replenishing repaired dNTP pool. Hsp31 members are widely recognized for metabolic and transcriptional reprogramming during the diauxic shift as they extensively tune the expression of necessary proteins for rapid acclimatization (Miller-Fleming et al., 2014). These observations may point to a unique regulatory mechanism by Hsp31 class of proteins similar to hDJ-1 in governing the chromatin landscape’s carbonyl biology (Zheng et al., 2019).

**Figure 8.**
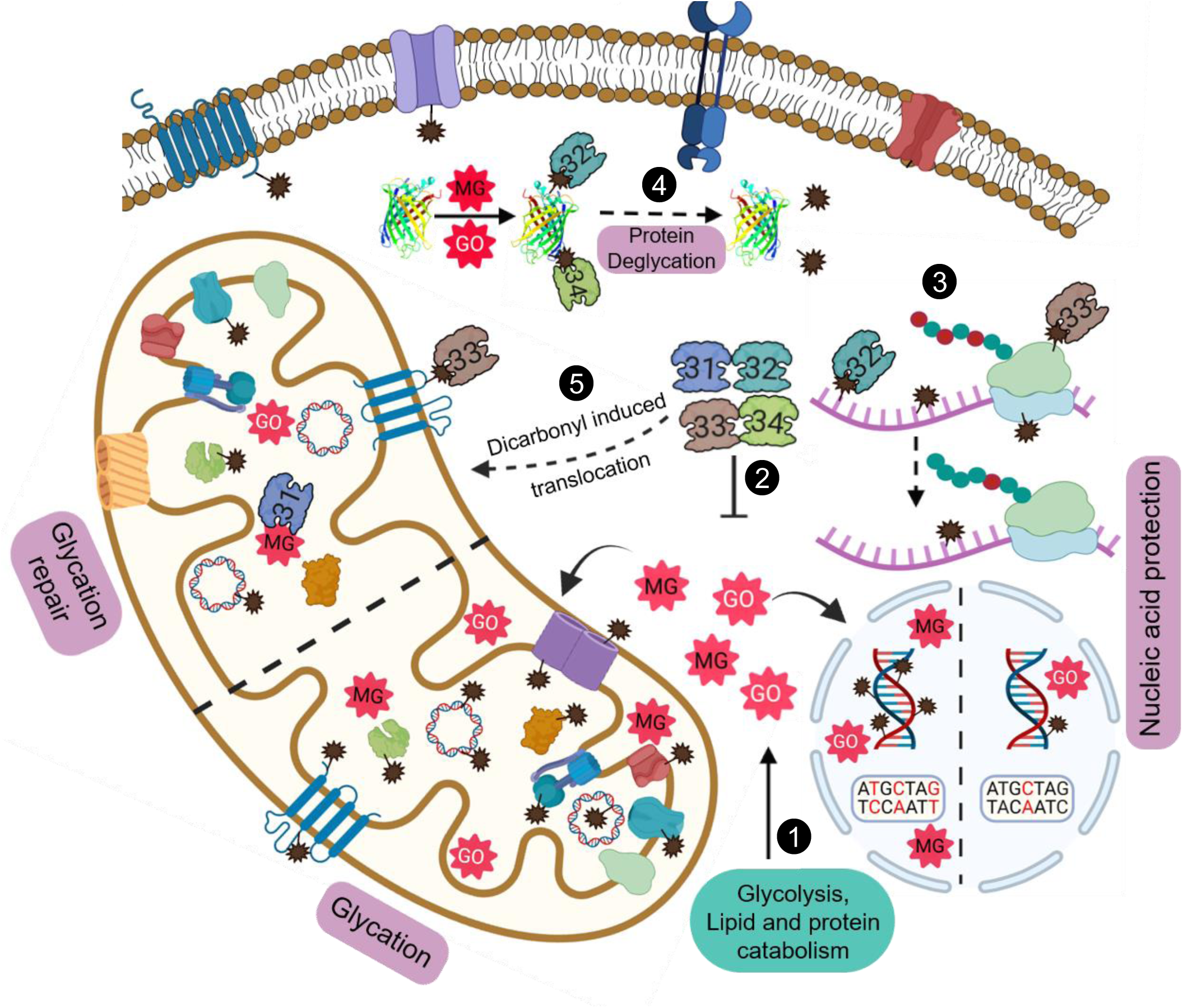
Model depicting the role of yeast DJ-1 orthologs in the maintenance and protection against carbonyls. Various metabolic pathways generate MG and GO that spontaneously glycate cytosol and organellar macromolecules **(1)**. Hsp31 paralogs scavenge excess endogenous RCS and regulate their levels **(2)**. The dual role of paralogs provides nucleic acid protection by efficient detoxification of carbonyls and reversal of damaged molecules **(3)**. Deglycation of proteins preserves the functional integrity of proteins **(4)**. Dicarbonyl stress-induced translocation of Hsp31 paralogs into mitochondria confers enhanced organellar protection by attenuating damage to mitochondrial biomolecules **(5)**.

Besides DNA, we found enriched glycation of RNA in Δ*Q* due to harmful levels of MG and GO. These modifications further disrupt the mRNA translation at the global scale, possibly through compromised binding or stalling of ribosomes. Significantly, yeast DJ-1 members extensively reverted the AGE modifications on RNA, suggesting their comprehensive role in translation regulation. The enhanced spectrum of mutations observed in various essential genes of Δ*Q* is a crucial indicator of a dedicated glycation repair pathway in *S. cerevisiae* (*Figure 8*). Interestingly, a recent report on *Arabidopsis thaliana* identified a novel glycation repair pathway similar to *S. cerevisiae*, involving AtDJ-1D in the repair of DNA and proteins (Prasad et al., 2022). Supporting the previous studies (Galligan et al., 2018; Nair et al., 2018; Richarme et al., 2017), we report yeast DJ-1 orthologs as a novel class of deglycases that repair RCS-mediated glycation of nucleic acids.

### Hsp31 paralogs regulate the proteome glycation homeostasis

Proteins are the most abundant biomolecules with considerably longer half-lives, making them highly vulnerable to multiple non-enzymatic covalent modifications. Under physiological conditions, glycation on arginine, lysine, and cysteine modulates the functional diversity of various proteins (Sun, Suttapitugsakul, Xiao, & Wu, 2019). We show that the absence of Hsp31 paralogs triggers severe glycation of the proteome, which may contribute to protein aggregation, misfolding, and cross-linking. Furthermore, under these circumstances, the conventional proteasomal degradation pathway fails to provide quality turnover resulting in AGE accumulation (Raupbach, Ott, Koenig, & Grune, 2020). As a compensatory response mechanism, amino acid biosynthesis and protein translation were upregulated in the single deletions of Hsp31 mini-family proteins (Miller-Fleming et al., 2014). This primarily indicates the biogenesis of functional proteins to promote cell survival during glycation stress. Therefore, scavenging excess endogenous RCS by the Hsp31 paralogs is imperative to evade persistent glycation and maintain balanced proteostasis. Besides, their robust deglycase activity repairs severely glycated proteins from different cellular compartments, including mitochondria. This enables cells to efficiently reutilize the cellular proteome pool (*Figure 8*). In contrast, human DJ-1 exhibited weaker methylglyoxalase and deglycase activity than yeast and plant counterparts (Mazza et al., 2022).

Due to its chemical properties and redox sensing ability, the catalytic cysteine in DJ-1 superfamily proteins holds critical importance during multi-stress responses (Wilson, 2011). In agreement with hDJ-1, C138A mutants of Hsp31 paralogs failed to repair macromolecules and detoxify RCS, suggesting a common multipurpose catalytic site.

### Yeast DJ-1 orthologs provide mitochondrial protection during carbonyl stress

MG and GO toxicity is one of the leading factors contributing to mitochondrial dysfunction in most neurodegenerative diseases (Wang, Xu, Musich, & Lin, 2019). Mechanistically, they disrupt the mitochondrial morphology and electron transport chain and elevate ROS levels. Although DJ-1 superfamily proteins display dynamic subcellular localization during the stress response, their physiological relevance in mitochondrial protection is not well studied (Junn, Jang, Zhao, Jeong, & Mouradian, 2009; Miller et al., 2003; Nair et al., 2018). In a previous report, we showed the maintenance of mitochondrial morphology by regulating the fusion-fission network by Hsp31 and Hsp34 (Bankapalli et al., 2020). Here, we provide direct evidence of mitochondrial maintenance by yeast DJ-1 orthologs, effectively through robust redistribution under glycation stress. Interestingly, translocation of Hsp31 members into mitochondria substantially lowers the glycation of mitochondrial proteome and DNA, resulting in persistent survival of mitochondria when challenged with RCS. Moreover, such redistribution of the paralogs could elevate the glycolate synthesis from glyoxal, a critical antioxidant that stabilizes mitochondrial membrane potential and re-establishes redox balance by producing GSH (Diez, Traikov, Schmeisser, Adhikari, & Kurzchalia, 2021; Toyoda et al., 2014). These findings highlight a possible neuroprotective function of PD-associated hDJ-1, involving the repair of damaged macromolecules and mitochondrial redistribution. Furthermore, such translocation into mitochondria can significantly revert mitochondrial dysfunction by attenuating glycation damage of mitochondrial DNA and proteins, thereby preventing the progression of PD.

In summary, our findings significantly contribute to understanding the diversified response by Hsp31 paralogs in the field of carbonyl biology. The repair pathway confers cytoprotection by preserving macromolecular and organellar integrity during chronic exposure to RCS. This shows an underlying mechanism of how hDJ-1 might provide neuroprotection, especially in the case of neurons that have intrinsically higher levels of RCS given the metabolic demand. Since Hsp31 paralogs are multi-stress responding proteins with new emerging functions, detailed investigations may further decipher the mechanistic insights on how they provide genome protection. Moreover, Hsp31 paralogs also translocate into stress bodies and P-granules, which are recently associated with neurodegenerative diseases (Dudman & Qi, 2020; Hallacli et al., 2022; Miller-Fleming et al., 2014). Therefore, it is critical to characterize the significance of their translocation as it would unravel a broader understanding of DJ-1 family members, specifically in hDJ-1, whose loss of function leads to familial PD.

## Materials And Methods

### Strains and plasmid constructions

The yeast strains used in this study are mentioned in *Supplementary file 1*. The details of the primers and plasmids utilized in this study are described in *Supplementary file 1*. Knockouts of Hsp31 paralogs were generated through homologous recombination in haploid yeast strain BY4741 (*MATα his3*Δ1 *leu2*Δ0 *met15*Δ0 *ura3*Δ0). Deletion of *hsp32* (Δ*32*), *hsp33*(Δ*33*), and *glo1*(Δ*glo1*), was performed by transforming suitable cassettes generated by primers P1-P4, constituting the *hphNT1* marker. Subsequently, the deletions were confirmed through PCR using primers P5-P8. The strains lacking hsp31 (Δ*31*) and hsp34 (Δ*34*) were previously generated in the lab. The double, triple (Δ*T*), and quadruple (Δ*Q*) deletion strains were generated by transforming DNA cassettes in Δ*31*Δ*34*. All the deletion strains were confirmed through PCR using primers P5-P8. RNR3 and Hsp31 paralogs expression was determined by genomically tagging at the C-terminus with heme agglutinin (HA) tag using primer P9-P11 and P12-P14, respectively, with *hphNT1* cassette as described above. Primers P15-P18 were used to tag RAD52 with GFP at the carboxy terminus in WT and Δ*Q*.

For overexpression studies, Hsp31 paralogs containing C-terminus heme agglutinin (HA) tag were cloned into pRS415 vector with GPD (glyceraldehyde-6-phosphate dehydrogenase) promotor and transformed into Δ*Q*. For protein expression and purification, Hsp31 family proteins were cloned into pRSF-Duet and purified with a previously published protocol (Bankapalli et al., 2015).

### Western blot analysis

Indicated respective strains grown till the mid-log phase (*A_600_*∼0.6) were treated with 10 mM MG or 15 mM GO for 12 h in the culture tubes. Later, the cell lysates were prepared by incubating with 10% Trichloro acetic acid solution at 4⁰C. Following acetone washes, 1X SDS dye (50 mM Tris-HCl, pH 6.8, 2% SDS, 0.1% bromophenol blue, 10% glycerol, and 100 mM b-mercaptoethanol) and acid-washed glass bead was added and vortexed thoroughly. Subsequently, the samples were heated at 92⁰C, and 30 µg of the sample was loaded. The lysates were resolved on 12% SDS-PAGE and transferred to the PVDF membrane. Subsequently, the membrane was blocked with 5% BSA in phosphate-buffered saline (PBST-Tween 0.05%). Later, the membrane was incubated with anti-CML or anti-MAGE antibody 1:1000 each overnight at 4⁰C. Following 3 washes with PBST, the membrane was incubated with the secondary antibody (1:10,000) at room temperature. Subsequently, the membrane was washed with PBST and exposed to luminol solutions (BIO-RAD). Equal loading of samples was confirmed through ponceau S staining of the membrane.

To determine the expression levels of Hsp31 paralogs in response to GO treatment, the WT strain containing genomic HA tag on the C-terminus of Hsp31 members was incubated in the absence or presence of 15 mM GO for 12 h. Subsequently, the cell lysates were resolved on 15% SDS-PAGE and probed with an anti-HA antibody. Similarly, WT and Δ*Q* were incubated without or with 0.01% MMS and 15 mM GO for 3 h to estimate the expression of RNR3. The fold change in all experiments was calculated by measuring the band intensities using Multi Gauge V3.0 software.

### Nucleic acid glycation and DNA dot blot analysis

Yeast strains grown till the mid-exponential phase were treated with 10 mM MG or 15 mM GO in the culture tubes for 12 h (Brown, 2001). Subsequently, genomic DNA was isolated using Wizard genomic DNA extraction kit (Promega, Cat No: A1120). For RNA isolation, cells were treated with 10 mM Tris pH 7.5, 1 mM EDTA, and 0.5% SDS (TES) buffer with hot acidic phenol for 1 h at 65⁰C. The resulting organic layer obtained after centrifugation was mixed with chloroform and centrifuged to acquire pure RNA. DNA and RNA were sheared through sonication and dotted (3 µg) on the nitrocellulose membrane (BIO-RAD, Cat No: 1620112). The membrane was baked at 80⁰C for 2 h and blocked-in phosphate-buffered saline (PBS-Tween 0.05%) with 5% BSA for 2 h at room temperature. Consequently, the membrane was probed overnight with an anti-CML or anti-MAGE antibody. Following the washes, the membrane was incubated with a secondary anti-mouse antibody. The membrane was exposed to luminol solutions. The membrane was stained with methylene blue to confirm equal loading of the samples.

### Polysome profiling

The experiment was performed using an established protocol (Shaffer & Rollins, 2020). Briefly, WT and Δ*Q* strains were grown till the mid-log phase and treated with 15 mM GO for 4 h at 30⁰C, and post-treatment cycloheximide was added to immobilize the ribosomes on mRNA. The cell lysates were prepared by adding lysis buffer (10 mM Tris pH 7.4, 100 mM NaCl, 30 mM MgCl_2_, 100 µg/ml cycloheximide, 10 U RiboLock, and 1 mM PMSF) to the pellet and vortexed with glass beads. The lysates were loaded on SW41 tubes containing a 10% to 50% sucrose density gradient, followed by ultra-centrifugation. Later, the fractions were analyzed using a polysome profiler (Piston gradient fractionator; BIOCOMP).

The area under the curve of polysome and monosome was calculated using Origin 8.0, and the ratio was plotted in Prism GraphPad 5.0.

### Phenotypic analysis and growth conditions

Individual yeast strains were grown till mid-log phase(*A_600_*∼0.6) in YPD (yeast extract-1%, peptone-2%, and dextrose-2%) or synthetic dropout (SD) lacking leucine amino acid (0.67% yeast nitrogen base without amino acids, 0.072% Leu dropout supplement and 2% dextrose). Consequently, cells were pelleted and serially diluted 10-fold until 10^-5^ dilution factor. Each dilution was spotted on the media plate without or with 15 mM GO. Parallelly, the cell pellet was treated with water or 10 mM MG for 6 h, serially diluted, and spotted on media plates. The plates were incubated at 30⁰C for 36 h and imaged. Phenotype under DNA damaging agents was determined by spotting cells from the mid-exponential phase onto YPD media plates containing 0.01% MMS (methyl methanesulfonate) or 150 mM HU (hydroxyurea). Also, the cells were spotted on YPD media plates containing 15 mM GO or 0.01% MMS and plates with both genotoxins. Images of spot assay were taken at 36 h.

### *In vitro* glyoxalase and methylglyoxalase activity

A previously established protocol was used to determine the activity (Bankapalli et al., 2015; C. Lee et al., 2016).To determine the enzyme activity, 5 µg of purified Hsp31 paralogs WT and C138A mutants were incubated with 0.5 mM GO or MG in HEPES KOH pH 7 buffer for 30 min at 30⁰C. Later, the reaction was terminated using 0.1% DNPH (2,4-dinitrophenylhydrazone), followed by adding 10% NaOH and incubating at 42⁰C for 10 min. The results were calorimetrically analyzed using Biospectrometer (Eppendorf), 570 nm for GO, and 530 nm for MG. The kinetic parameters were determined by incubating Hsp31 members with varying GO concentrations, and the readings were noted at different time points. The experiments were performed in three biological replicates and triplicates. Subsequently, the values were plotted on Prism GraphPad 5.0 to estimate the *K_m_* and *V_max_*.

### *In vitro* deglycation assay of DNA and Proteins

The deglycation of DNA and protein was performed using previously published protocols with minor modifications (Richarme et al., 2017). For DNA deglycation experiments, 500 µM dNTPs or 150 µM forward and reverse primers of yeast *sod1* (P19-P20) were incubated without or with GO/MG (2 mM) in HEPES KOH buffer pH 7 at 30⁰C for 2 h. Before supplementing the reactions with Hsp31 paralogs, the glycation mixtures were first diluted (6-fold) to minimize the free GO or MG levels. Post glycation reaction, 5 µg of purified Hsp31 paralogs and BSA were supplemented and incubated for 3 h at 30⁰C. The deglycase activity of Hsp31 paralogs was determined by subjecting the samples to a polymerase chain reaction (PCR) to amplify the *sod1* gene in the presence of Phusion enzyme high fidelity DNA polymerase.

To measure the protein deglycation activity, 2 µg of purified human Sod1 and Lysozyme were treated without or in the presence of 2 mM of GO/MG in HEPES KOH buffer pH 7 at 30⁰C for 2 h (Prasad et al., 2022). Subsequently, the samples were diluted 6-fold to minimize GO or MG residual levels. Post glycation of protein samples, the reaction mixture was supplemented with 5 µg of purified Hsp31 paralogs and incubated for 3 h for deglycation. The deglycation status of proteins was determined by resolving the samples on 15% SDS-PAGE, followed by western analysis and immunodetected with anti-CML or anti-MAGE antibodies.

### Genetic mutation and sequencing

Respective strains were grown till the mid-log phase and treated with 15 mM GO or 10 mM MG in the culture flasks for 24 h at 30⁰C. Total DNA was isolated as described above, and PCR amplified for *sod1*, *rad14*, and *cox2* ORFs using primers P19-P24. Subsequently, the amplicons were sequenced by the Sangar method, and the raw reads were aligned with the WT sequences. The number of mutations was determined using the multi-sequence alignment tool Clustal Omega and the data was plotted on GraphPad Prism 5.0.

### Microscopic analysis

To analyse the formation of RAD52 GFP foci, WT and Δ*Q* cells were grown until the exponential phase and treated with 0.03% MMS and 15 mM GO for 1 h. Subsequently, the *A_600_* = 0.5 OD cells were harvested and washed with 1X PBS before spreading over 2% agarose pads.

The localization of yeast DJ-1 members was performed by genomically tagging with GFP at the carboxy terminus using primers P12-P14. Consequently, the cells were transformed with a pRS415_TEF_ vector expressing MTS-mCherry to decorate mitochondria (Bankapalli et al., 2015). The transformed strains grown until the mid-exponential phase were harvested (*A_600_* = 0.5) and incubated with water or 15 mM GO for 2 h. Parallelly, the cells were treated with water or 10 mM MG for 2 h and harvested. The cell pellet was washed with 1X PBS and processed for imaging. All the images were acquired in confocal microscopy (Olympus FV3000) with a 10 µm scale bar.

### Chemicals and antibodies

Chemicals used in the study are Glyoxal (Sigma,128465), Methylglyoxal (Sigma, M0252), Methyl Methanesulfonate (TCI, M0369), Hydroxyurea (Sigma, H8627), Lysozyme (Sigma, 4403), and nitrocellulose membrane (BIO-RAD,1620112). Antibodies used in the study are anti-CML (Carboxymethyl lysine; MAB3247 R&D Systems, Minneapolis, MN, USA), anti-MAGE (Argpyrimidine specific; ab243074 Abcam), anti-HA antibody (GT4810, Sigma), secondary mouse (GE-bioscience).

### Miscellaneous

Mitochondria were isolated from cells treated with 15 mM GO for 16 h using a previously published protocol (Matta, Kumar, & D’Silva, 2020).

### Statistical analysis

Glycation intensities of DNA and proteins were measured through quantitation of complete dots or lanes, respectively, in Multi Gauge V3.0 software. The values were normalized with respect to WT in all experiments and plotted in GraphPad Prism 5.0 software. Error bars represent the standard deviation derived from three biological replicates. One-way ANOVA with Dunnett’s multiple comparisons test was used to determine significance analysis, comparing multiple columns against the WT. Asterisks used in the figures represent the following significance values: *, *p*≤ 0.05; **, *p*≤0.01; ***, *p*≤ 0.001; ****, *p* ≤ 0.0001; NS, not significant.

## Acknowledgments

We thank Prof. Umesh Varshney (Department of Microbiology and Cell Biology, Indian Institute of Science, Bangalore, India) for the utilization of polysome profile machine. We acknowledge the confocal microscope facility (Department of Biochemistry, Indian Institute of Science, Bangalore, India). Prof. Patrick D’Silva acknowledges financial support from DST-SERB grant (CRG/2018/001988), Department of Biotechnology (DBT-IISC Partnership Program Phase-II, No. BT/PR27952/IN/22/212/2018), and DST-FIST Program-Phase III (No. SR/FST/LSII045/2016-G). Gautam Susarla and Priyanka Kataria acknowledges the fellowship from Indian Institute of Science-Bangalore, India. Amrita Kundu acknowledges the fellowship from Department of Biotechnology-Research associate (DBT-RA), India.

## Additional information

### Funding

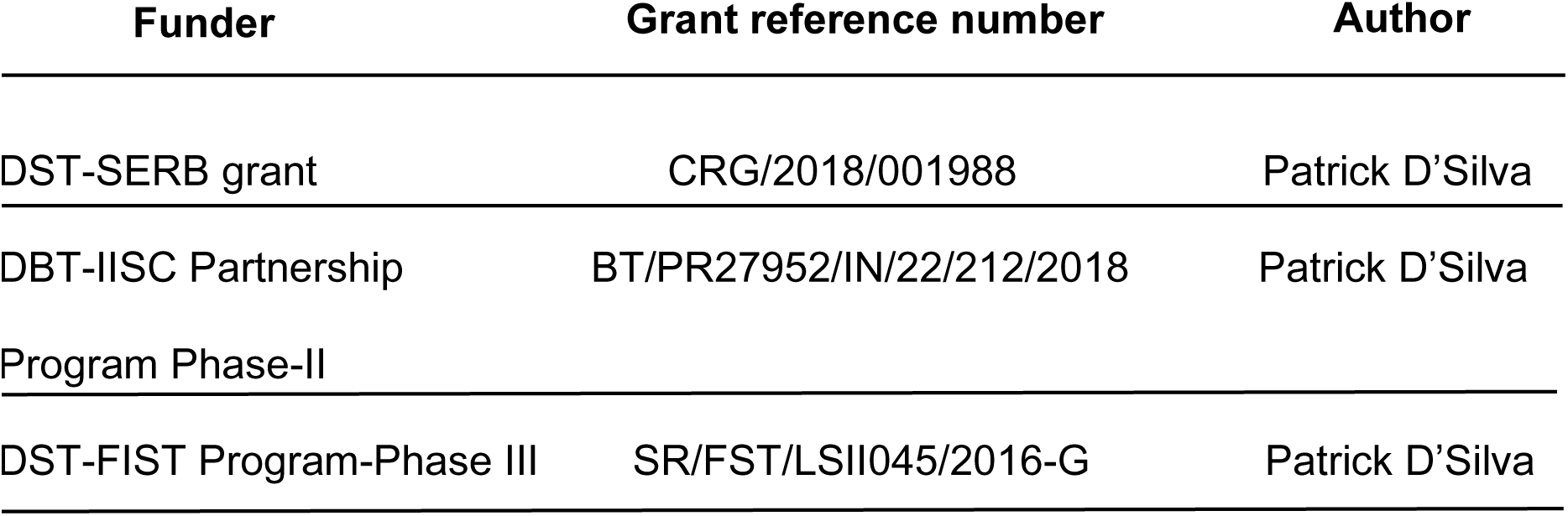

### Author contributions

Gautam Susarla, Conceptualization, Data curation, Formal analysis, Investigation, Methodology, Validation, Visualization, Writing-Original draft, Writing-review and editing; Priyanka Kataria, Amrita Kundu, Data curation, Methodology, Validation; Patrick D’Silva, Conceptualization, Formal analysis, Fund acquisition, Project administration, Supervision, Validation, Writing-review and editing.

### Additional files

- Source data files contains both the original image data and Image data before trimming used in Figures and Figure supplements.
- Supplementary file 1. List of strains, primers, and plasmids used in this study.
- MDAR checklist

### Data Availability

All data generated or analysed during this study are included in the manuscript and supporting files; Source data files have been provided.

## Figure legends

**Figure 1-figure supplement 1.**
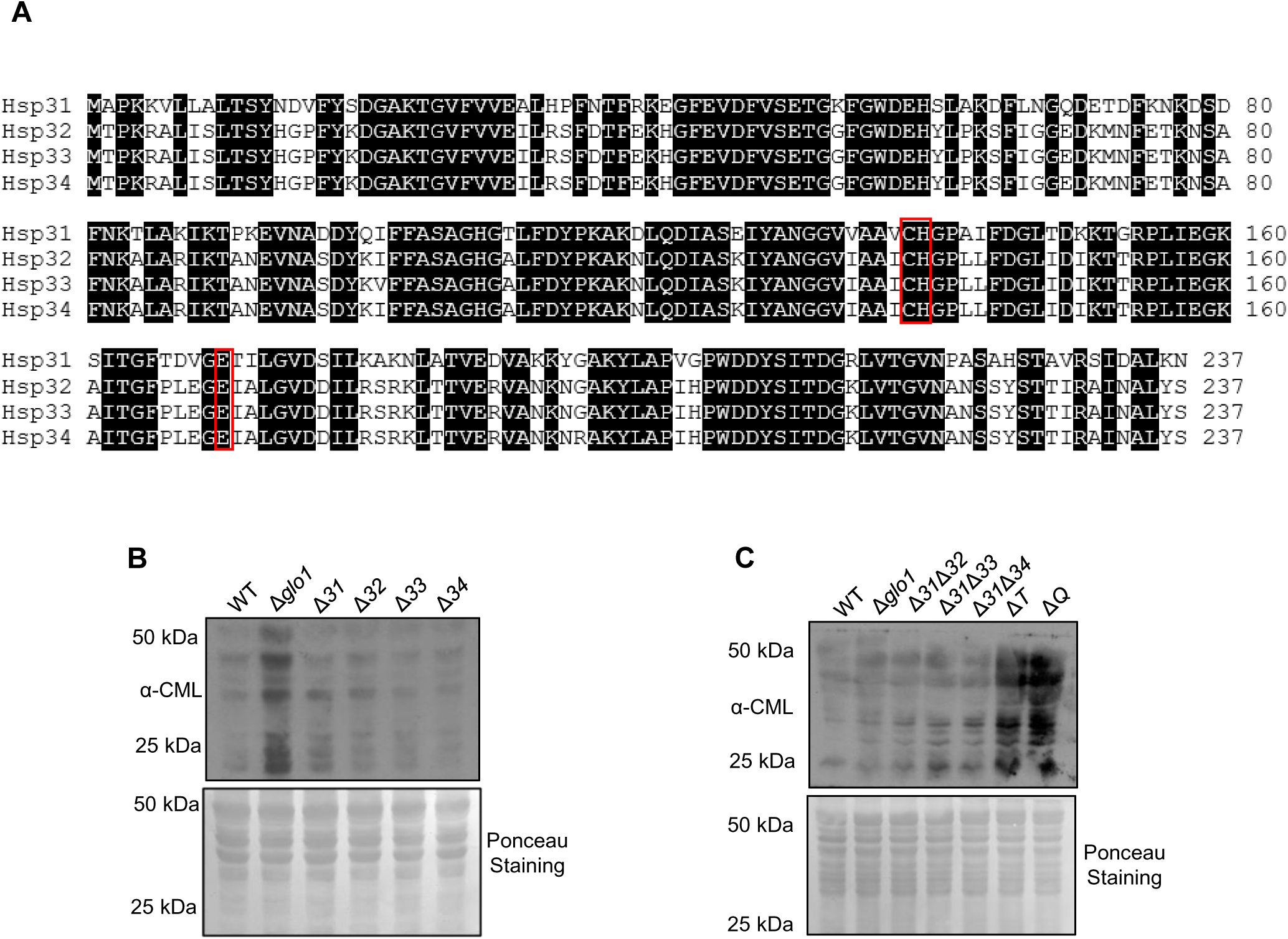
Amino acid sequence alignment of Hsp31 paralogs and protein glycation profile in the absence of GO stress. **(A)** Amino acid sequences of Hsp31 paralogs were aligned using the online tool Clustal Omega. Black shade indicates identical amino acids. Red boxes indicate amino acids of the catalytic triad. **(B, C)** Respective strains were grown till the stationary phase and subsequently lysed. The protein glycation levels were estimated by immunodetection using anti-CML antibody. **Figure 1—figure supplement 1—source data 1.** Source data for *Figure 1—figure supplement 1B*. **Figure 1—figure supplement 1—source data 2**. Source data for *Figure 1—figure supplement 1C*.

**Figure 2-figure supplement 1.**
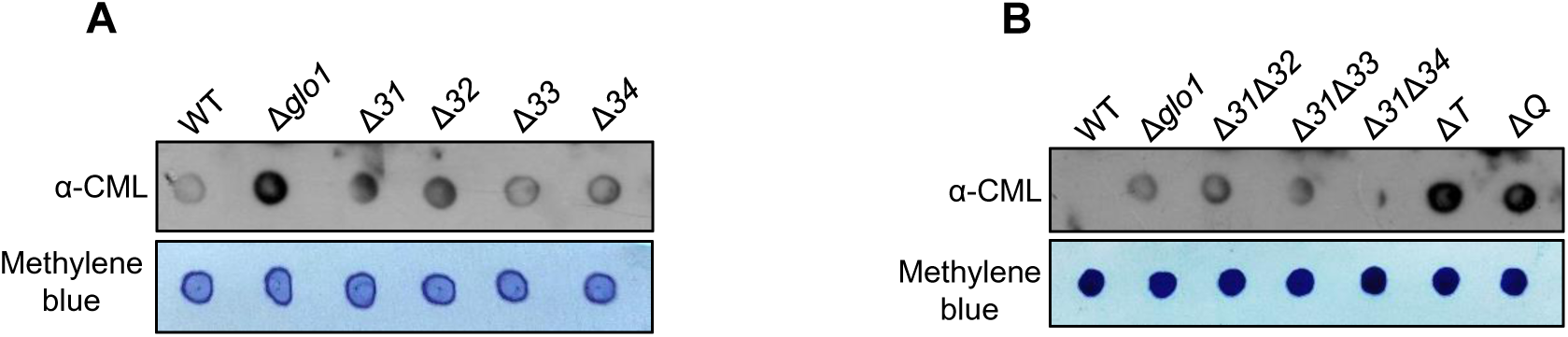
Glycation profile of DNA in the absence of GO stress. **(A, B)** DNA extracted from stationary phase cells was dotted (3 µg) on nitrocellulose membrane and probed with anti-CML to determine the glycation status **Figure 2—figure supplement 1—source data 1.** Source data for *Figure 2—figure supplement 2A*. **Figure 2— figure supplement 1—source data 2**. Source data for *Figure 2—figure supplement 2B*.

**Figure 3-figure supplement 1.**
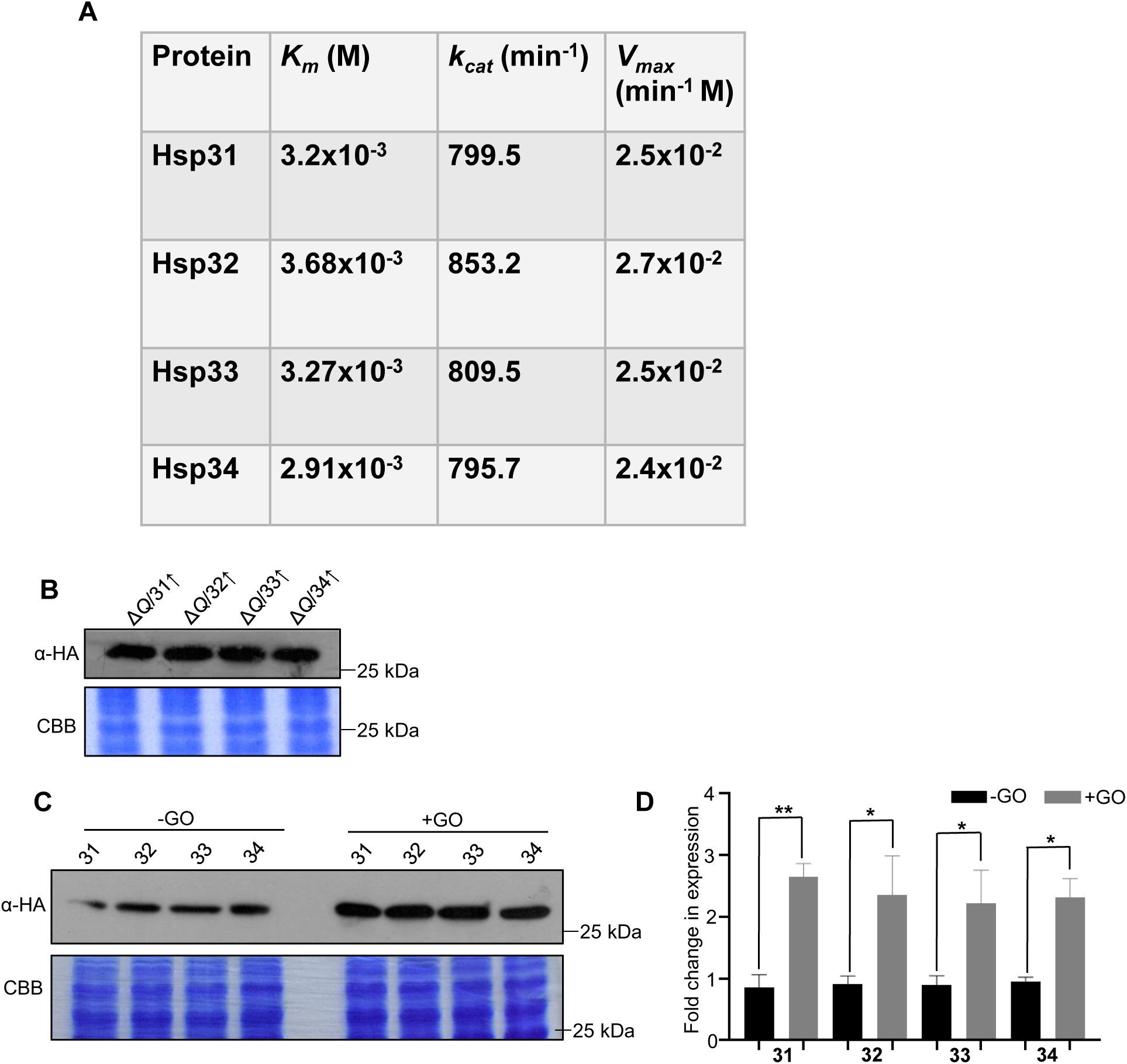
Enzyme kinetics and expression of yeast DJ-1 orthologs under GO stress. **(A)** GO was incubated at various concentrations with 5 µg Hsp31 paralogs, and the reaction was terminated at different time intervals. The corresponding readings were plotted on Prism GraphPad 5.0 and calculated for *K_m_*, *k_cat_*, and *V_max_*. **(B)** Hsp31 members expressed using the plasmid pRS415_GPD_ in Δ*Q*, confirmed with western analysis. **(C, D)** GO induced expression of Hsp31 paralogs. Cells expressing Hsp31 members with genomic tagged HA were incubated without or with 15 mM GO for 12 h, and the expression was determined by immunoblotting with an anti-HA antibody. Gel stained with Coomassie brilliant blue (CBB) was used as a loading control. The fold change in the expression was calculated using Multi-Gauge V3.0 and plotted in Prism GraphPad 5.0. **Figure 3—figure supplement 1—source data 1.** Source data for *Figure 3—figure supplement 3B*. **Figure 3—figure supplement 1—source data 2.** Source data for *Figure 3—figure supplement 3C*. **Figure 3—figure supplement 1—source data 3**. Source data for *Figure 3—figure supplement 3D*.

**Figure 5-figure Supplement 1.**
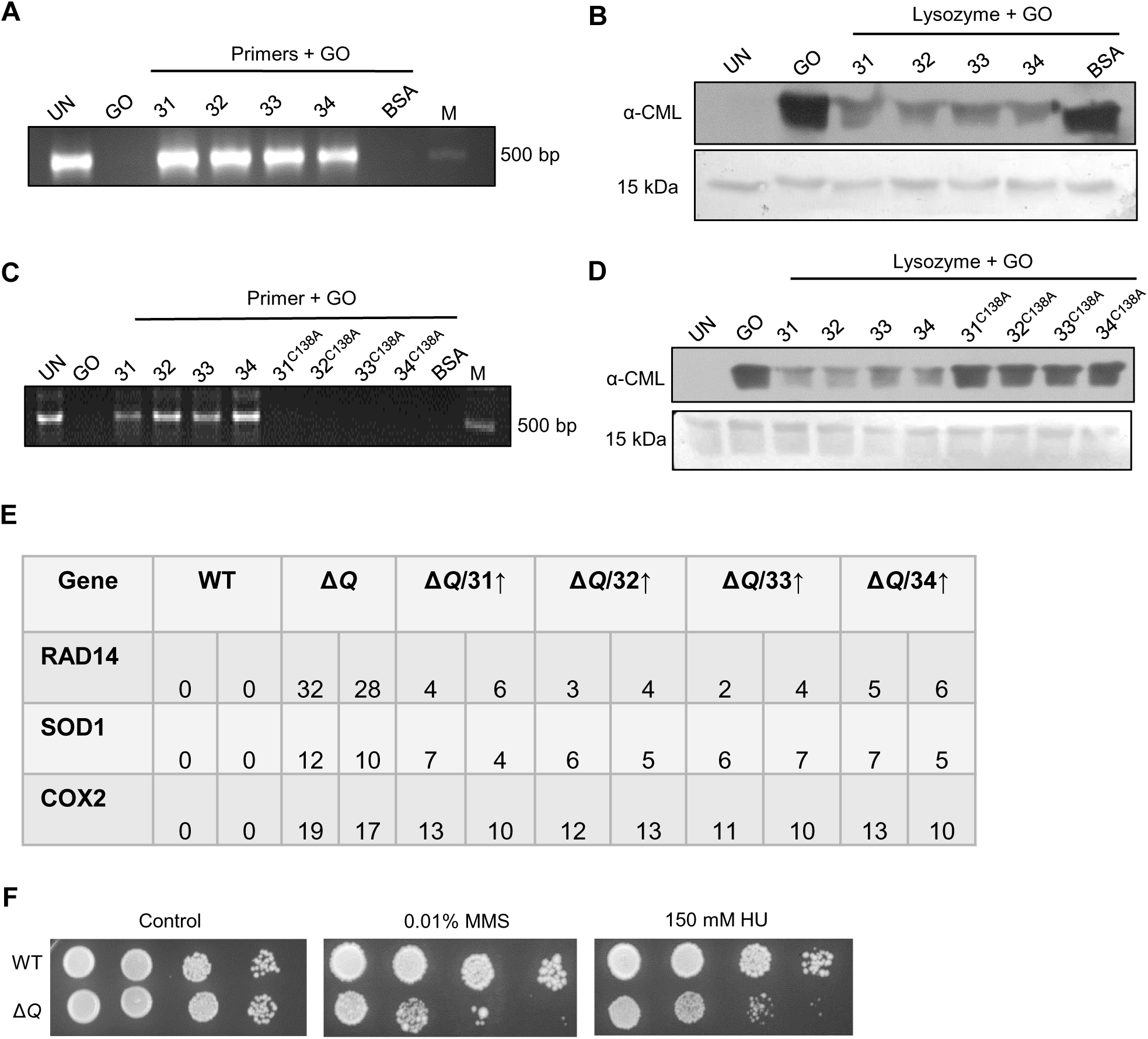
GO-associated macromolecular deglycation, mutation frequency profile, and genotoxic damage. **(A-D)** DNA oligonucleotides (150 µM) and Lysozyme (2 µg) were incubated for 2 h in the absence and presence of 2 mM GO, followed by deglycation with Hsp31 paralogs and C138A mutants for 3 h. **(E)** Respective genes were PCR amplified from cells treated with 15 mM GO for 16 h and Sanger sequenced. The number of genetic mutations from various genes is represented here. **(F)** Phenotypic analysis under DNA damaging agents. The cells harvested from the mid-log phase were spotted on plates containing 0.01% MMS or 150 mM HU. Plates were incubated at 30⁰C and imaged at 36 h. **Figure 5—figure supplement 1—source data 1.** Source data for *Figure 5—figure supplement 5A*. **Figure 5—figure supplement 1—source data 2.** Source data for *Figure 5—figure supplement 5B*. **Figure 5—figure supplement 1—source data 3.** Source data for *Figure 5—figure supplement 5C*. **Figure 5—figure supplement 1—source data 4.** Source data for *Figure 5—figure supplement 5D*.

**Figure 6-figure supplement 1.**
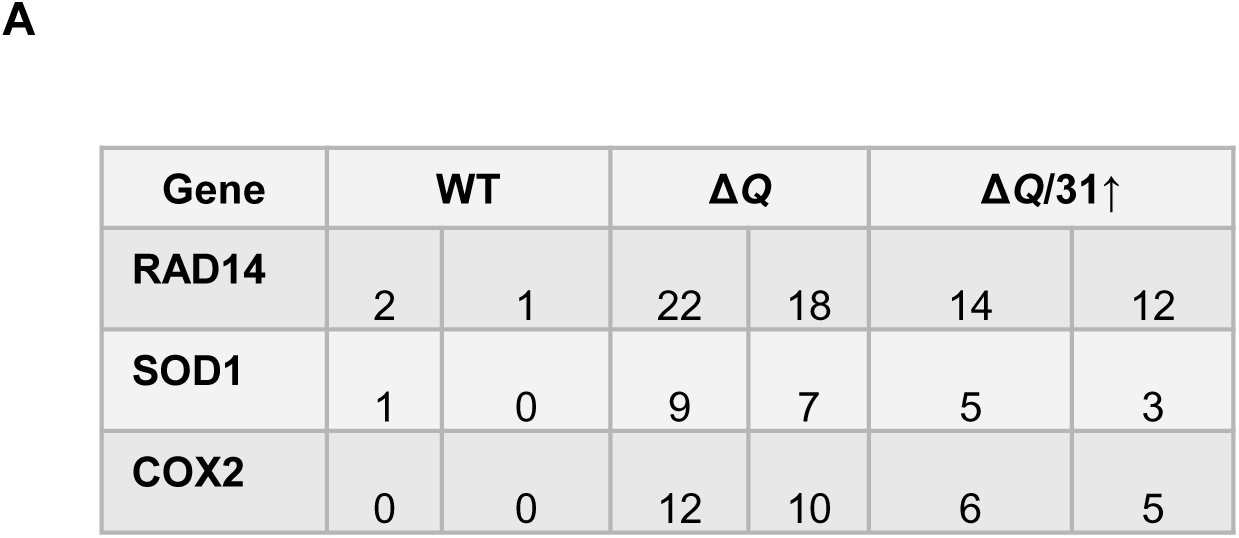
Mutation frequency in the presence of MG. **(A)** Individual genes, as indicated, were PCR amplified from various strains incubated with 10 mM MG for 16 h and Sanger sequenced. The number of genetic mutations was determined and represented here.

**Figure 7-figure supplement 1.**
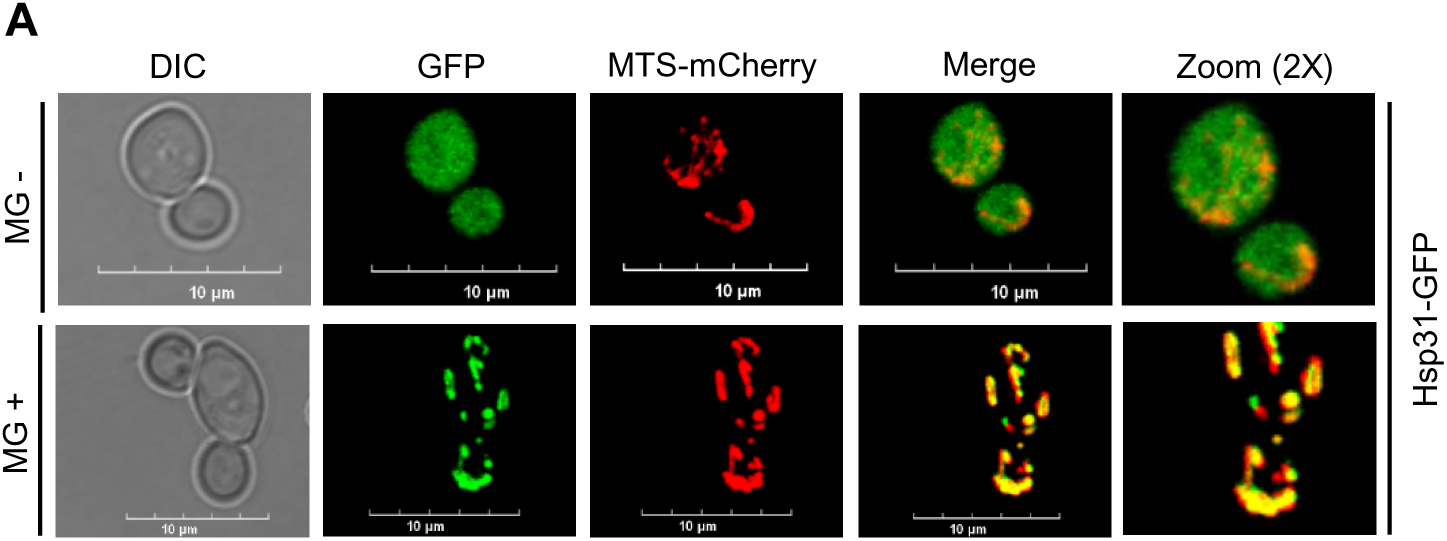
MG induced translocation of Hsp31 into mitochondria. **(A)** WT strains expressing C-terminus GFP-tagged Hsp31 and MTS-mCherry were treated without (-MG) or with 10 mM MG (+MG) for 3 h. Consequently, images were captured in a confocal microscope (Olympus FV3000), and represented with 10 µm scale.

